# Inhibition of FAM19A5 restores synaptic loss and improves cognitive function in mouse models of Alzheimer’s disease

**DOI:** 10.1101/2023.11.22.568357

**Authors:** Han-Byul Kim, Sangjin Yoo, Hoyun Kwak, Shi-Xun Ma, Ryunhee Kim, Minhyeok Lee, Nui Ha, Soonil Pyo, Soon-Gu Kwon, Eun-Ho Cho, Sang-Myeong Lee, Juwon Jang, Wonkyum Kim, Hae-Chul Park, Minkyung Baek, Yosub Park, Ji-Young Park, Jin-Woo Park, Sun Wook Hwang, Jong-Ik Hwang, Jae Young Seong

## Abstract

**Introduction:** Alzheimer’s disease (AD) is characterized by the dysregulation of synaptic balance, with progressive loss of synapses outpacing formation, ultimately leading to cognitive decline. However, the lack of effective strategies for restoring lost synapses poses a major barrier to improving clinical outcomes.

**Methods:** We developed NS101, a monoclonal antibody targeting FAM19A5, a brain-secreted protein. Its preclinical efficacy in restoring synapses and cognition was evaluated using APP/PS1 and P301S mice. The clinical safety and target engagement of NS101 were examined in human participants.

**Results:** FAM19A5 binds to LRRC4B, a postsynaptic adhesion molecule, leading to synapse reduction. Blocking this interaction with NS101 normalized the rate of synapse elimination in AD mice. This synaptic rebalancing restored the number and function of synapses, resulting in improved cognition. Systemically administered NS101 facilitated the transport of brain FAM19A5 into the bloodstream.

**Discussion:** Targeting FAM19A5 may hold clinical promise for treating AD by restoring synaptic balance.

## 1. BACKGROUND

FAM19A5, also known as TAFA5, is a secretory protein primarily detected in neurons of the central and peripheral nervous systems^1^. Studies using FAM19A5 promoter-driven LacZ expression combined with qRT[PCR analysis have shown that FAM19A5 is highly expressed in the central nervous system rather than in peripheral tissues such as adipose tissues^2,3^. These findings align with single-cell/nuclear RNA-seq data from humans and mice, indicating predominant expression of FAM19A5 in neurons, moderate expression in astrocytes and oligodendrocyte precursor cells, and minimal expression in oligodendrocytes and microglia^4–7^. Tracing the expression pattern of FAM19A5 in FAM19A5-lacZ knock-in mice revealed its presence in neural precursor cells during the early stages of embryonic development, suggesting that it plays a pivotal role in neural growth and circuit establishment^8^. Recent knockout studies have further highlighted the crucial role of FAM19A5 in the later stages of synaptogenesis and pruning processes, potentially in shaping synaptic connectivity and related brain functions^9,10^. Moreover, genome-wide association studies have linked FAM19A5 with various neurological disorders, including Alzheimer’s disease (AD), attention deficit hyperactivity disorder, and autism, highlighting its significance in the pathophysiology of these conditions^11,12^.

The amino acid sequence of FAM19A5 remains highly conserved across vertebrates, including humans, but the divergence from the FAM19A1-4 family is significant, indicating that FAM19A5 has a distinct function in the brain^13,14^. While FAM19A1-4 are known to interact with neurexin, a presynaptic adhesion molecule that participates in a wide range of neurological processes^15^, FAM19A5 is suggested to act as a ligand for different synaptic receptors. This action may be involved in synapse modification, thereby impacting neural function. However, the interaction of FAM19A5 with its unknown receptor and its downstream effects on synapse regulation in key brain regions, such as the cortex and hippocampus, remain elusive.

AD is a neurodegenerative disorder characterized by the progressive loss of synapses, manifested as an imbalance between synapse formation and elimination, leading to cognitive decline. While there are several therapeutic agents approved for treating AD, they offer only limited symptomatic relief and do not slow the progression of the disease^16^. Recently, antibody-based immunotherapy against amyloid beta (Aβ) has attracted significant attention following its approval by the US FDA^17–19^. However, synapse loss that continues after Aβ clearance may limit the efficacy of immunotherapy for cognitive improvement^19–21^. A novel approach for restoring lost synapses could be a breakthrough, as synapse loss, a fundamental hallmark of AD, contributes to cognitive deficits^22^. Understanding the mechanisms underlying synapse loss in AD may facilitate the development of more potent treatments^23–26^.

Here, we propose an antibody therapy targeting FAM19A5 that is capable of restoring synapses and cognitive function despite the presence of amyloid or tau aggregates. We found that FAM19A5 binds to LRRC4B, a postsynaptic adhesion molecule, through the FAM19A5 binding (FB) domain. This binding inhibited the interaction between LRRC4B and PTPRF, a presynaptic adhesion molecule, potentially leading to synapse elimination. Conversely, inhibiting FAM19A5 with a competitive protein derived from the FB domain attenuated synapse elimination, resulting in an increased number of synapses in primary neurons. These findings suggest that FAM19A5 acts as a physiological modulator of synaptic balance by facilitating controlled synapse elimination.

However, in AD, the influx of toxic pathological aggregates disrupts the delicate equilibrium of synapse formation and elimination, tilting the balance toward excessive elimination. Given this dysregulation, we hypothesized that the inhibition of physiological synaptolytic factors such as FAM19A5 could rebalance the tilted equilibrium. NS101, an anti-FAM19A5 antibody, effectively neutralized FAM19A5 and normalized the synaptic balance in the brains of tauopathy and amyloidopathy mice. This, in turn, resulted in improved cognitive performance. Systemically administered NS101 was successfully delivered to the brain, facilitating dose-dependent transport of brain FAM19A5 to the bloodstream in rodents and humans. This finding raises the potential for noninvasive clinical applications.

## 2. METHODS

### 2.1 Human samples

Pooled normal human cerebrospinal fluid (CSF) was obtained from Innovative Research, Inc. (IPLA-CSFP) and used for co-immunoprecipitation (Co-IP) experiments to determine the association between FAM19A5 and LRRC4B. Individual human CSF and plasma samples were obtained from BIOIVT & ELEVATING SCIENCE and were used to measure age-dependent FAM19A5 levels in CSF and the basal level of FAM19A5 in plasma, respectively. Additionally, individual human CSF samples were also used to determine total tau and phospho-tau levels.

### 2.2 Co-immunoprecipitation

HEK293 cells were transfected with reagents from the Neon^TM^ Transfection Kit (Invitrogen) according to the manufacturer’s instructions. The primer sets used for Co-IP of the construct are listed in Table S1. For the general Co-IP experiments, after transfection, the cells were treated with FAM19A5 for 30 min. The cells were washed twice with cold phosphate-buffered saline (PBS) and resuspended in lysis buffer containing 20 mM Tris-HCl (pH 7.4), 150 nM NaCl, 0.5% NP40, and a protease and phosphatase inhibitor cocktail (Thermo Scientific). The supernatants were isolated by centrifugation at 12,000 × g at 4°C for 30 min and then mixed with 30 μl of Protein G Dynabeads (Invitrogen) that had been preincubated with antibodies. The mixtures were incubated at 4°C overnight in a rotating mixer. The beads were spun down and washed three times with 20 mM Tris-HCl (pH 7.4), 300 mM NaCl, and 0.5% NP40 in washing buffer. After adding 30 μl of 2× sample buffer containing a reducing agent to the beads and boiling at 100°C for 10 min, the proteins were separated via SDS[PAGE.

For human CSF Co-IP, pooled human CSFs (Innovative Research) were mixed with 0.5% NP40 and 50 μl of Pierce Protein A/G Plus Agarose (Thermo Scientific) that had been preincubated with 10 μg of anti-human IgG, N-A5-Ab, and C-A5-Ab. The mixtures were rotated at 4°C overnight and washed three times. The protein complexes in the beads were resolved by SDS[PAGE.

For Co-IP, endogenous proteins from the mouse cerebral cortex were dissected and lysed in RIPA buffer (Thermo Scientific) supplemented with 25 mM Tris-HCl, 150 mM NaCl, 1% NP-40, 1% sodium deoxycholate, 0.1% SDS, and a protease and phosphatase inhibitor cocktail (Thermo Scientific). The tissue was then homogenized and subjected to sonication to ensure complete lysis. To remove insoluble material, the lysate was centrifuged at 16,000 × g for 30 minutes at 4°C. Subsequently, 1 mg of cortex lysate supernatant was incubated with 30 μl of Protein G Dynabeads (Invitrogen) preincubated with the appropriate antibodies (the information for antibodies is in Table S2): anti-FAM19A5 (S-A5-Ab), anti-LRRC4B (Alomone labs), and anti-PSD95 (Invitrogen). The subsequent steps were conducted as described for the Co-IP procedure performed with HEK293 cells.

### 2.3 ELISA

To evaluate both total and phosphorylated tau levels, ELISA kits, such as hTAU Ag (Fujirebio) and PHOSPHO-TAU (Fujirebio), were used according to the manufacturer’s instructions.

For the measurement of FAM19A5 in biofluids and tissues, we coated 96-well microplates with LRRC4B (453-576) protein or S-A5-Ab, which was diluted in 50 mM carbonate buffer (pH 9.6) to achieve a final concentration of 1 µg/ml. The plates were sealed and incubated overnight at 4°C. The following day, we washed the plates twice with 300 µL of washing buffer (PBS with 0.05% Tween 20) per well using a microplate washer (Tecan) and gently tapped them on a paper towel to remove any remaining solution. Subsequently, 200 μl of blocking buffer was added to each well, and the plates were sealed and incubated at 37°C for 1 hour. After two additional washes, 100 μl of both the standard solution and the samples were added to each well. The plates were sealed and incubated at room temperature for 90 minutes. Following another five washes, 100 μl of HRP-conjugated C-A5-Ab, diluted in blocking buffer (PBS with 1% BSA and 0.05% Tween 20) to a final concentration of 0.2 μg/ml, was added to each well and incubated at 37°C for 1 hour. Then, 100 μl of TMB solution (Thermo Scientific) was added to each well, and the plates were incubated at room temperature for 20 minutes. To stop the colorimetric reaction, 100 μl of 1 N sulfuric acid was used, and the optical density (OD) of each well was determined using a microplate reader (Molecular device) at 450 nm.

To measure CSF NS101, we used a pair of rabbit anti-human IgG heavy chain antibodies (Invitrogen) and HRP-conjugated goat α-human IgG kappa light chain antibodies (Thermo Scientific).

To assess the binding affinity between FAM19A5 and the LRRC4 family, we utilized various LRRC4/4B/4C deletion proteins, including LRRC4 (39-527), LRRC4B (36-576), LRRC4B (453-576), LRRC4B (484-576), LRRC4B (498-576), and LRRC4C (45-527), as capture agents. We used both the WT and MT FAM19A5 proteins, which included FAM19A5 (R58A, R59A), FAM19A5 (R125A, K127A), and FAM19A5 (R58A, R59A, R125A, K127A), for binding analysis. Detection was carried out using HRP-conjugated C-A5-Ab.

To examine the inhibition of FAM19A5-LRRC4B binding, we first coated plates with the LRRC4B (453-576) protein. Subsequently, we added the test sample and the FAM19A5 protein. The degree of binding inhibition was determined by detecting FAM19A5 using an HPR-conjugated C-A5-Ab. For the PTPRF-LRRC4B binding inhibition test, the LRRC4B (36-576) protein was coated on the plate, followed by the addition of the test sample and PTPRF protein. Binding inhibition was calculated by detecting PTPRF-hFc using an anti-human IgG Fc antibody.

The amino acid sequences used to elucidate the key residues in the FB domain of LRRC4B and to identify the epitopes of the FAM19A5 antibodies are listed in the Tables S3, S4, and S5.

### 2.4 Counting of dendritic spines

Dendritic spines were imaged from layers II/III of the prefrontal cortex and the CA1 of the hippocampus using a 63x oil-immersion lens with 3x digital zoom. Segments of 30 μm long basal dendrites were sampled midway between the soma and the distal termination (medial). A total of 160 images were obtained from the brain sections (80 μm thick at 0.5 μm intervals). Maximum intensity projection (MIP) images were obtained from a total of 160 images and processed with ImageJ software (NIH). Five MIP images per subject were selected for quantification. The dendritic spine counter in ImageJ was used for dendritic spine counting and spine classification. At least 15 dendrites per subject were measured, and spines were classified as mushroom, stubby, thin, or filopodium using the criteria^27,28^.

### 2.5 Two-photon live spine imaging

Thirty-two female P301S mice (B6;C3-Tg (Prnp-MAPT*P301S) PS19Vle/J; JAX Strain #008169), as well as sixteen female B6C3F1/J (JAX Strain #100010) mice, were anesthetized with ketamine/xylazine and subjected to cranial window implantation^29^. Before surgery, the mice received carprofen (subcutaneous, 5 mg/kg) to relieve pain during and after surgery and dexamethasone (subcutaneous, 2 μg/g) to reduce inflammation. The cranial window was inserted over the somatosensory cortex at the following coordinates: AP −1.8 and ML −2.0 from the bregma. A dental drill (HP4–917, Foredom) was used to remove a round (d = 4 mm) piece of skull, and the hole in the bone was covered with a round cover glass (d = 5 mm; Electron Microscopy Sciences, 72296–05). A helicopter-shaped head plate (model 1; Neurotar) was placed over the cover glass and fixed to the skull surface with dental cement (Rapid Repair; Dentsply) mixed with cyanoacrylate glue (Loctite 401; Henkel). An AAV viral vector encoding GFP under the control of the synapsin promoter (AAV9-synapsin-GFP, Signagen, SL100845-Std) was injected intracortically during surgery (300 nanoliters, titer 1:10, depth of injection 800-900 micrometers from the cortical surface) at the following coordinates: AP −1.8 and ML −2.0 from the bregma to fluorescently labeled neurons. Among the 48 mice that underwent surgery, 30 mice with the most transparent cranial windows were selected for imaging in the study 3 weeks after implantation. Four weeks after the surgery, the mice were habituated to the head fixation during the 4 days of training. Therefore, imaging started after 4 weeks of postsurgical recovery. Identical brain ROIs were imaged for 66 days at 7 fixed time points (days 1, 5, 9, 13, 17, 38, and 66).

Seven-month-old P301S mice were imaged with an FV1200MPE two-photon microscope (Olympus, Japan) with a 25X water immersion 1.05 NA objective specially designed for in vivo two-photon imaging. A MaiTai Broad Band DeepSee laser tuned to 900 nm was used for excitation. Emission light was collected using a bandpass filter (515–560 nm). For in vivo imaging sessions, awake animals were head-fixed under a two-photon microscope using the Mobile Home Cage device (Neurotar, Finland). One week prior to the start of imaging, the animals were habituated to the head fixation conditions for four days.

After image acquisition, the images were processed and analyzed using ImageJ (NIH). First, shifts between time points in the x, y, and z directions were compensated using Neurotar’s scripts in ImageJ. Spine turnover analysis was performed by a scientist completely blinded to the identity of the treatment groups. Individual spines were tracked manually by visual identification in 3D stacks of images collected at all time points. Protrusions of at least 5 pixels from the dendritic shaft were considered candidate spines. After identification of a candidate spine, the data analyst examined 10 adjacent slices in both directions to determine other explanations for the protrusion. If no other explanation was found, the spine was counted in the analysis. At every time point, each spine was assigned a specific code, either 1 (present) or 0 (absent). If, in some instances, the image quality was not sufficiently sharp for spine identification at a specific time point, the spine was assigned a code -1 and excluded from further analysis.

### 2.6 Quantification and statistical analysis

All the statistical analyses were performed using GraphPad Prism 8 with 95% confidence. All the data are presented as the mean ± SEM. All the details of the statistical tests and results are reported in the figure legends.

## 3 RESULTS

### 3.1 FAM19A5 interacts with LRRC4B

We hypothesized that FAM19A5, as a synapse modification factor, may interact with a synaptic adhesion molecule other than neurexin^15^ to regulate synaptic balance (Figure 1A). To explore potential binding molecules for FAM19A5, we investigated RNA expression patterns from 168 types of neurons in the mouse brain scRNA-seq database^4^, comparing their correlation with *fam19a5* to narrow down candidates. We found that the expression of *lrrc4b*, a postsynapse adhesion molecule, was highly correlated with that of *fam19a5*, suggesting a possible interaction in the synaptic cleft (Figure 1B and Figure S1). Given their similar expression patterns and their secretion into the cerebrospinal fluid (CSF)^30–32^, we expected that FAM19A5 and LRRC4B would exist in a complex form in the CSF. To determine the potential interaction between these two proteins, we conducted Co-IP with human CSF. We selectively captured FAM19A5 from human CSF using monoclonal antibodies against each N-and C-terminal region of FAM19A5 and separated them by size through western blotting to identify bound proteins. Indeed, LRRC4B was co-immunoprecipitated with the antibody that binds to the C-terminal region of FAM19A5, suggesting that FAM19A5 and LRRC4B may form a complex in the brain (Figure 1C).

**FIGURE 1.**
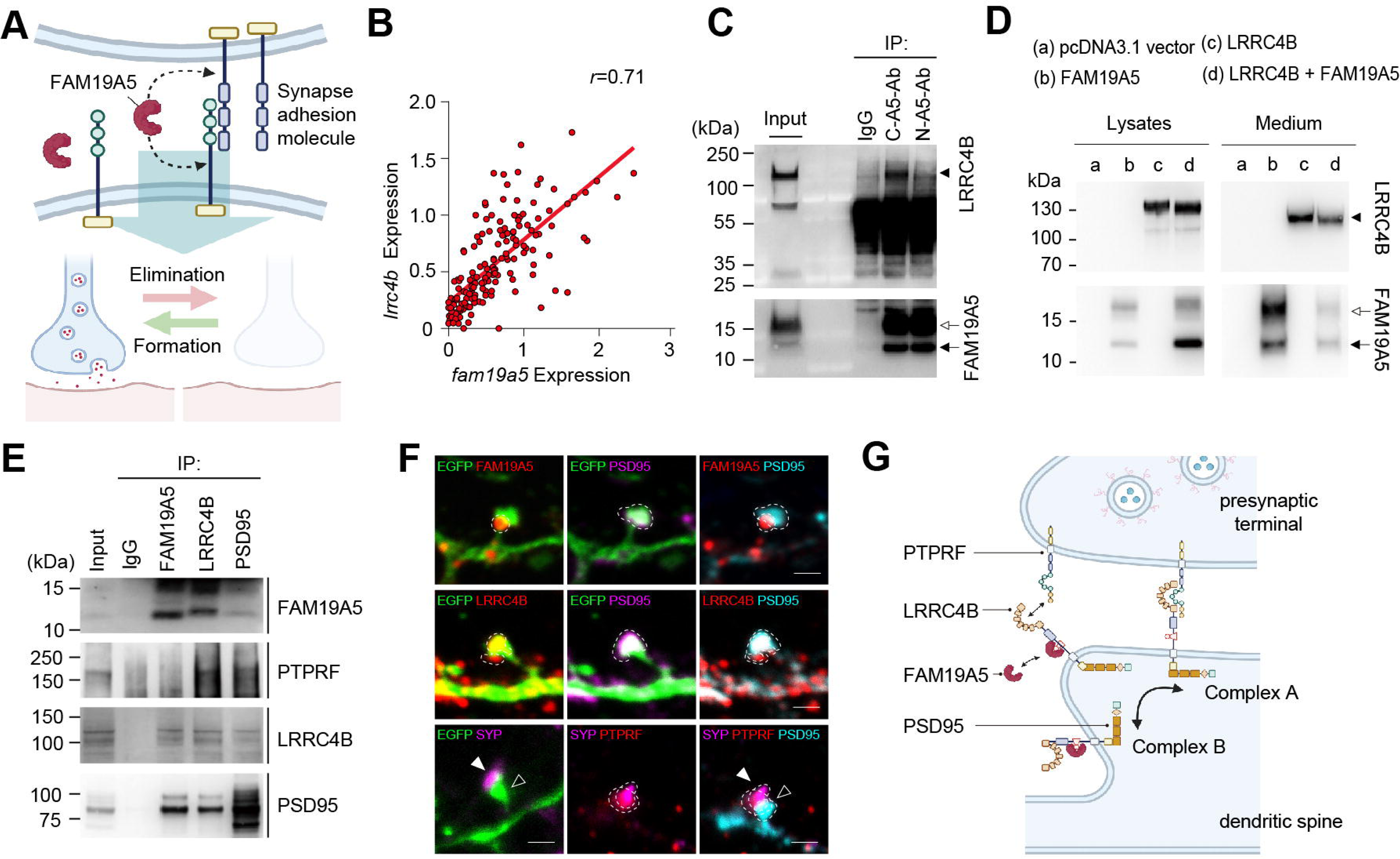
FAM19A5 binds to LRRC4B. (A) Illustration of the potential mechanisms of synapse regulation. FAM19A5 binding to adhesion molecules could induce either synapse elimination or formation. (B) Correlations between the RNA transcript levels of *lrrc4b* and *fam19a5* in 168 neuron types. Correlation coefficient; *r*= 0.71, *P* < 0.0001. (C) Co-IP of human CSF using anti-FAM19A5 antibodies (C-A5-Ab or N-A5-Ab). Solid arrowhead: LRRC4B, open arrow: glycosylated FAM19A5, solid arrow: nonglycosylated FAM19A5. (D) Immunoblots of LRRC4B and FAM19A5 in culture lysate and medium from HEK293 cells coexpressing LRRC4B and FAM19A5. Solid arrowhead: LRRC4B, open arrow: glycosylated FAM19A5, solid arrow: nonglycosylated FAM19A5. (E) Co-IP using lysates prepared from the mouse cerebral cortex to investigate the interaction between FAM19A5, PTPRF, LRRC4B and PSD95. (F) Representative image of a dendritic spine from a cultured hippocampal neuron at 13 DIV showing the localization of the synaptic proteins FAM19A5 (red), PTPRF (red), SYP (magenta, presynaptic marker), LRRC4B (red/cyan, postsynaptic adhesion molecule), and PSD95 (magenta/cyan, postsynaptic marker). The locations of synaptic proteins are represented by dotted lines. Solid arrowhead: presynapse, open arrowhead: postsynapse. Scale bar, 1 μm. (G) Illustration of the potential regulation of synaptic adhesion molecules by FAM19A5. The formation of complexes A and B, consisting of PSD95-LRRC4B-PTPRF and PSD95-LRRC4B-FAM19A5, respectively, is regulated by FAM19A5.

Western blot and Co-IP results revealed two distinct forms of FAM19A5 of different sizes that bind to LRRC4B, which can be attributed to the presence or absence of posttranslational modifications such as glycosylation. Indeed, glycosylated FAM19A5 proteins in mouse and human CSF samples were deglycosylated by the PNGase F enzyme (Figure S2A). Additionally, we confirmed that both glycosylated and deglycosylated FAM19A5 bind to LRRC4B with similar affinities: the EC50 values for naïve and nonglycosylated FAM19A5 were 107 and 90 pM, respectively (Figure S2B), suggesting minimal functional differences between the forms.

The FAM19A5-LRRC4B interaction was further confirmed by immunoblotting assays with coexpressing cells and medium. LRRC4B detected in the medium is a secreted form cleaved at the juxta-membrane region of full-length LRRC4B^33^. This cleavage resulted in a lower molecular weight than that of LRRC4B in the cell lysate. We observed that upon coexpressing LRRC4B, the concentration of secreted FAM19A5 in the medium decreased, while its concentration in the lysates markedly increased (Figure 1D). These results demonstrate that LRRC4B is a specific binding molecule for FAM19A5 that effectively captures FAM19A5 and prevents its release into the medium.

We then conducted Co-IP using lysates prepared from the mouse cerebral cortex to investigate whether FAM19A5 interacts with LRRC4B under physiological conditions. We also explored the involvement of PSD95, a postsynaptic scaffold protein, and PTPRF, a presynaptic cell adhesion protein, in this interaction because they are known to interact with LRRC4B at synapses^34,35^. Our Co-IP results revealed that FAM19A5 forms a complex with LRRC4B and PSD95 but not with PTPRF. Interestingly, LRRC4B also forms a complex with PTPRF, independent of the FAM19A5-LRRC4B-PSD95 complex. PSD95 forms complexes with both LRRC4B and PTPRF, with FAM19A5 weakly bound to this complex (Figure 1E). These findings suggest the formation of two distinct complexes at the synapse: one consisting of FAM19A5-LRRC4B-PSD95 and another without FAM19A5, consisting of PTPRF-LRRC4B-PSD95. These results suggest that FAM19A5 may be a key molecule regulating the physical connection between the presynapse and postsynapse.

Immunocytochemistry experiments on cultured hippocampal neurons further corroborated the Co-IP findings. These experiments revealed that FAM19A5 colocalizes with PSD95 and LRRC4B at postsynaptic sites. Notably, FAM19A5 was primarily observed in the spine neck and only weakly in the spine head, where PSD95 was predominantly concentrated. LRRC4B exhibited a broader distribution, being found in both the spine neck and head. Furthermore, PTPRF largely colocalized with the presynaptic marker synaptophysin (SYP) in contact with postsynaptic spines, suggesting that PTPRF may help bridge pre- and postsynaptic sites through interactions with LRRC4B (Figure 1F). These findings, together with the Co-IP results, suggest that the dissociation of FAM19A5 from the LRRC4B complex is necessary for the relocation of LRRC4B to the spine head to interact with presynaptic proteins. Alternatively, the association of FAM19A5 with LRRC4B might promote the disengagement of LRRC4B from presynaptic proteins, facilitating its movement from the spine head to the neck and potentially aiding in synapse disassembly (Figure 1G).

### 3.2 FAM19A5 binds to the FB domain of LRRC4B

To determine which domain of LRRC4B interacts with FAM19A5, we generated a series of LRRC4B fragments with different domains (Figure 2A) and performed Co-IP assays. HEK293 cells expressing FLAG-tagged LRRC4B deletion constructs were treated with rcFAM19A5 proteins for 30 min. Cell lysates were immunoprecipitated with anti-FLAG M2 affinity gel, and immunoprecipitated proteins were immunoblotted with anti-FLAG and N-A5-Ab. We found that all constructs containing the 484–498 aa sequence of LRRC4B were able to bind to FAM19A5, whereas deletion of the 484–498 aa sequence abolished FAM19A5 binding (Figure 2B, S3A). Using ELISA, we then tested this interaction using purified proteins, confirming both the binding affinity and interaction. In line with the Co-IP results, FAM19A5 exhibited high affinity for LRRC4B fragments within the amino acid range of 484-498 (EC_50_ values were 68.5 pM for the sequence 453-576 and 112.1 pM for 484-576). In contrast, no binding was evident without the sequence (Figure 2C). These results demonstrate that the LRRC4B sequence in the range of residues 484-498 acts as a FAM19A5 binding domain, termed the FB domain. Surface plasmon resonance (SPR) measurements revealed a pronounced binding affinity between FAM19A5 and LRRC4B (453-576), with an equilibrium dissociation constant (K_D_) of 32 pM (Figure S3B). This affinity notably stands out from that reported for other synaptic adhesion molecules^36^.

**FIGURE 2.**
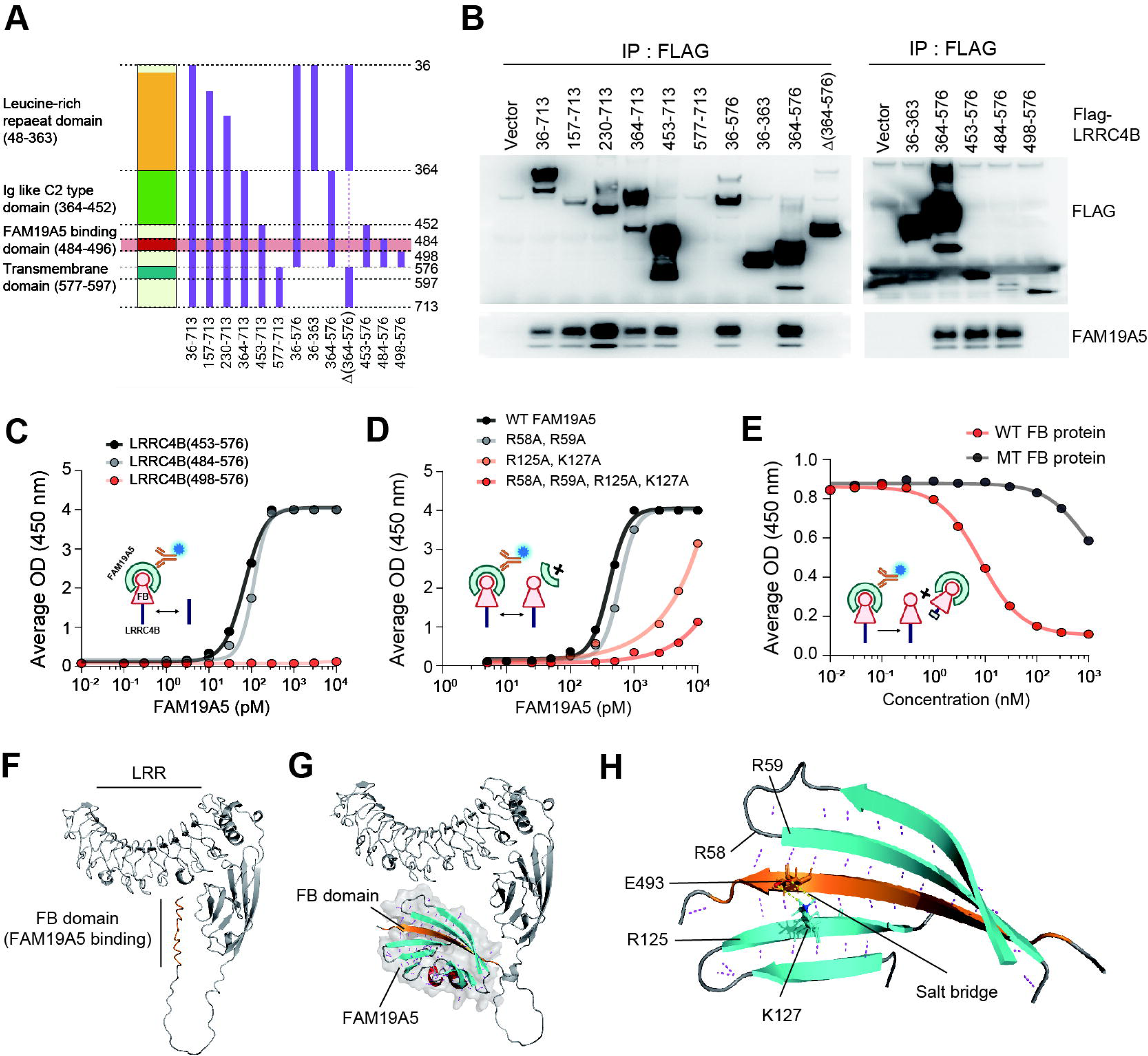
Characterization of FAM19A5-LRRC4B binding. (A) Domain architecture of the LRRC4B deletion constructs. (B) Co-IP of interactions between FAM19A5 and LRRC4B deletion constructs. (C) The binding affinity of LRRC4B for FAM19A5 was determined via ELISA (inserted, captured by LRRC4B-hFc, detected by HRP-conjugated C-A5-Ab). The EC_50_ of LRRC4B(453-576)-hFc = 68.5 pM, and the EC_50_ of LRRC4B(484-576)-hFc = 112.1 pM. (D) Binding affinity of wild-type (WT) or mutant (MT) FAM19A5 proteins (R58A, R59A; R125A, K127A; R58A, R59A, R125A, K127A) to LRRC4B measured via ELISA (inserted, captured by His-TEV LRRC4B, detected by HRP-conjugated C-A5-Ab). The EC_50_ of WT FAM19A5 = 409.5 pM, and the EC_50_ of MT FAM19A5 (R58A, R59A) = 594.8 pM. (E) Inhibition of FAM19A5-LRRC4B binding by WT LRRC4B(453–576)-hFc or MT LRRC4B(453–576, T488A, T489A)-hFc FB-containing proteins determined via ELISA (inserted, captured by LRRC4B(36-576)-hFc, detected by HRP-conjugated C-A5-Ab). The IC_50_ of the WT FB-containing protein was 8.1 nM. (F) *In silico* model of LRRC4B, including the leucine-rich repeat (LRR), immunoglobulin C (IgC), and FB domains (orange). (G) *In silico* model of the LRRC4B-FAM19A5 complex. The FAM19A5 structures (gray surface) are depicted as α-helices (red) and β-strands (cyan). (H) Model showing the salt bridge between the side chains of E493 (orange) and K127 (cyan). Hydrogen bonds are represented by dashed lines. The predicted key residues of FAM19A5 involved in binding to the FB domain (484-498, orange) are R58, R59, R125, and K127.

To further elucidate the essential residues within the FB domain, we investigated the binding free energy and compared its changes after the substitution of each residue with Ala. These results indicated that all the residues, except Thr494 and Leu495, played vital roles in binding (Figure S3C), which was further supported by the ELISA results (Figure S3D). We also determined the key residues of FAM19A5 involved in LRRC4B binding. Residues Arg58, Arg59, Arg125, and Lys127 of FAM19A5 were predicted to be significant binding residues (Figure S3E). Subsequent ELISAs demonstrated that Arg125 and Lys127 were significant for LRRC4B binding, while Arg58 and Arg59 contributed moderately to the interaction (Figure 2D). Interestingly, the amino acid sequence of the FB domain is highly conserved across vertebrate species, including humans (Figure S3F), implying crucial function of the FAM19A5-LRRC4B complex in the brain. To further validate the FAM19A5-FB interaction, we measured FAM19A5-LRRC4B binding in the presence of WT or MT FB-containing protein (LRRC4B[453–576, T488A, T489A]-hFc), which do not bind to FAM19A5. We observed that the WT FB-containing protein effectively inhibited the binding of FAM19A5 to LRRC4B in a dose-dependent manner, whereas the MT FB-containing protein had minimal impact (Figure 2E). The competition experiments using WT and MT FB-containing proteins strongly support the specific interaction between the FB domain of LRRC4B and FAM19A5.

Using AlphaFold2 and RoseTTAFold^37–39^, we performed 3D modeling of the FAM19A5-LRRC4B complex to delve into their structural and molecular interactions. Complex modeling showed that these proteins interact through the key residues of FAM19A5 and the FB domain of LRRC4B, supporting the results of the biochemical analyses shown in Figure 2A, B. In particular, the sequence of FB, which exhibited a disordered structure prior to binding with FAM19A5, was transformed into a β-strand. The β-strand then interacted with the β-strands of FAM19A5, primarily via hydrogen bonds (Figure 2F, G). Furthermore, predictions indicated the formation of a salt bridge between LRRC4B Glu493 and FAM19A5 Lys127, one of the pivotal residues for binding. This interaction likely further stabilized the complex structure (Figure 2H).

### 3.3 FAM19A5 binding to LRRC4B leads to the dissociation of synaptic contacts

Mouse primary hippocampal neurons endogenously expressed high levels of FAM19A5, LRRC4B, and PTPRF transcripts since their early stage of development (Figure S4). To investigate the role of FAM19A5 in synapse formation or elimination, we overexpressed FAM19A5 in hippocampal neurons. To visualize synaptic contacts, we cotransfected the neurons with plasmids containing either eGFP alone or eGFP along with FAM19A5. Subsequently, the neurons were immunostained with SYP to confirm the presence of presynaptic terminals (SYP-positive puncta) in contact with the dendritic spines of the eGFP-positive neurons. Using these images, we quantified spine density, particularly the density of spines in contact with SYP. Overexpression of FAM19A5 significantly reduced the density of spines, particularly those in contact with SYP, compared to neurons transfected with eGFP alone (Figure 3A, B).

**FIGURE 3.**
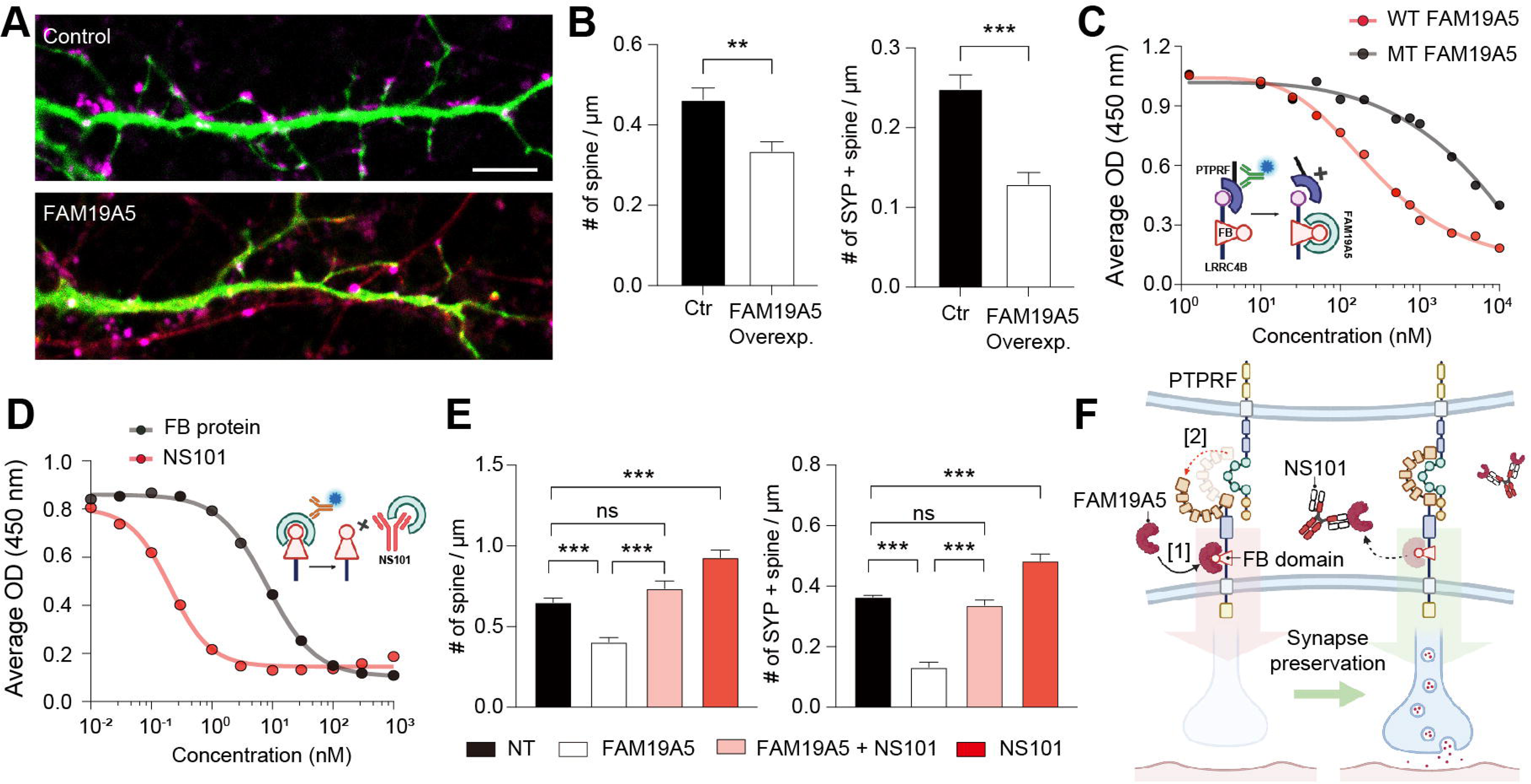
FAM19A5 binding to LRRC4B leads to the dissociation of synaptic contacts. (A) Representative image of dendritic spines on cultured hippocampal neurons transfected with either the eGFP control (green) or FAM19A5 (red). SYP (magenta), a presynaptic marker, is in contact with the spine, indicating possible synapse assembly. (B) Quantification of the number of spines and spines in contact with SYP (n = 7 per group). Data are presented as the mean ± SEM. Unpaired t-test was used to calculate *P* values. \*\**P* < 0.01, \*\*\**P* < 0.001. (C) Dissociation of the LRRC4B-PTPRF complex in the presence of WT FAM19A5 or MT FAM19A5 [R58A, R59A, R125A, K127A] determined via ELISA (inserted, captured by His-TEV LRRC4B, detected by HRP-conjugated anti-human IgG Fc). The IC_50_ of WT FAM19A5 = 281 nM. (D) Inhibition of FAM19A5-LRRC4B binding by NS101 or FB-containing proteins determined via ELISA (inserted, captured by LRRC4B(36-576)-hFc, detected by HRP-conjugated C-A5-Ab). The IC_50_ of NS101 was 205.0 pM, and the IC_50_ of the FB protein was 8.1 nM. (E) Quantification of the number of spines and spines in contact with SYP (n = 7 per group) in hippocampal neurons treated with NT, FAM19A5 (50 nM), or NS101 (50 nM) or cotreated with FAM19A5 and NS101 (50 nM). Data are presented as the mean ± SEM. One-way ANOVA followed by Tukey’s multiple comparison test was used to calculate *P* values. \*\*\**P* < 0.001. (F) Schematic illustration of how NS101 prevents synapse loss by inhibiting FAM19A5.

Given that LRRC4B forms a transsynaptic connection with PTPRF within the synaptic cleft^35^, we postulated that the binding of FAM19A5 to LRRC4B may induce a conformational change in the LRRC4B-PTPRF complex, resulting in synapse modification. To test this hypothesis, we performed ELISAs to quantitatively evaluate LRRC4B-PTPRF binding. We observed changes in binding upon the introduction of either wild-type (WT) or mutant (MT) FAM19A5 [R58A, R59A, R125A, K127A], which lacks binding affinity for the FB domain of LRRC4B. WT FAM19A5 significantly decreased LRRC4B-PTPRF binding in a dose-dependent manner, whereas MT FAM19A5 only partially inhibited this binding (Figure 3C, IC_50_ of WT FAM19A5: 281 nM). This result demonstrated that FAM19A5 can dissociate the LRRC4B-PTPRF complex.

Building upon this discovery, we hypothesized that conversely, detaching FAM19A5 bound to LRRC4B could promote synapse formation. For safe and specific targeting of FAM19A5 in the brain, we developed a monoclonal antibody against FAM19A5 called NS101. Among the FAM19A family members, NS101 exhibited specific binding to FAM19A5 (Figure S5A). SPR measurements showed that NS101 has a strong affinity for FAM19A5, with K_D_ =111 pM (Figure S5B). ELISAs using FAM19A5 fragments demonstrated that the epitopes for NS101 were Arg52, Pro57, Arg58 and Arg59 (Figure S5C). Notably, Arg58 and Arg59 are key residues of FAM19A5 for its binding to LRRC4B, indicating that NS101 can inhibit FAM19A5-LRRC4B complex formation by competitively binding to these key residues. Owing to this high affinity, NS101 was able to dissociate the FAM19A5-LRRC4B complex more effectively than the synthetic FB-containing protein (Figure 3D, IC_50_ for NS101: 0.2 nM vs. IC_50_ for FB-containing protein: 8 nM).

Next, we hypothesized that blocking FAM19A5 with NS101 could inhibit its binding to LRRC4B, thereby preserving synaptic connections. To validate this hypothesis, we exposed primary hippocampal neurons to FAM19A5 and/or NS101 and evaluated spine density, as shown in Figure 3A and B. Treatment of neurons with FAM19A5 significantly decreased spine density. However, when NS101 was administered together with FAM19A5, NS101 significantly blocked the effect of FAM19A5. Furthermore, treatment with NS101 alone significantly increased spine density (Figure 3E). Similarly, treatment of hippocampal neurons with WT FB-containing protein, but not with MT FB-containing protein, significantly increased the intensity and colocalization of SYP and PSD95 compared to those in the control group (Figure S6A, B).

Collectively, these findings suggest that FAM19A5, by specifically binding to LRRC4B, disrupts the LRRC4B-PTPRF complex within synapses, contributing to synapse elimination. However, removing FAM19A5 bound to LRRC4B using NS101 can preserve the complex and trigger the reassociation of LRRC4B-PTPRF, thereby restoring synapses (Figure 3F).

### 3.4 NS101 enters the brain and transports brain FAM19A5 into the peripheral circulation

Building upon the established role of FAM19A5 as a physiological synaptolytic factor, we hypothesized that amyloid plaques and tau tangles in AD act as pathological drivers of excessive synapse elimination, tilting the balance toward detrimental synapse loss. Our recent observation showed that reducing FAM19A5 levels in the AD brain by crossbreeding FAM19A5LacZ+/-mice with APP/PS1 mice extended the lifespan compared to that of APP/PS1 mice, partially supporting this hypothesis^40^. Therefore, for the treatment of AD, we targeted FAM19A5 to restore synaptic balance rather than removing pathological aggregates through the monoclonal FAM19A5 antibody NS101.

Before we proceed to AD mice, we first examined whether intravenously (IV) administered NS101 can enter the brain and bind to FAM19A5. To evaluate the pharmacokinetics of NS101, we monitored its concentration in rat blood following a single IV administration. NS101 was gradually excreted from the body over a span of 28 days, with a half-life of 7.5 days for a dose of 10 mg/kg and 11.7 days for a dose of 50 mg/kg (Figure 4A). The area under the curve (AUC) revealed a dose-dependent increase in systemic exposure to NS101 (10,336.3 hour·µg/mL for a dose of 10 mg/kg and 54,415.0 hour·µg/mL for a dose of 50 mg/kg). Similar to other clinical antibodies^41^, prolonged systemic exposure to NS101 facilitated its delivery to the brain across the blood-brain barrier (BBB). In the mouse brain, NS101 reached its peak concentration 30 hours after a 10 mg/kg IV injection. Subsequently, NS101 levels declined slowly over 28 days, with a half-life of 17.3 days (Figure 4B). Based on the 28-day AUC, the plasma-brain antibody ratio indicated that approximately 3% of the plasma NS101 reached the brain, which was comparable to the clinical antibody for Aβ^42^. NS101 in the brain was either excreted directly to the blood vessels or transported via the CSF. Following a single IV injection, NS101 reached its peak concentration in the rat CSF at 36 hours, exhibiting a dose-dependent increase (Figure 4C). The 28-day plasma-to-CSF antibody ratio indicates that approximately 0.06% of the plasma NS101 was transported to the CSF.

**FIGURE 4.**
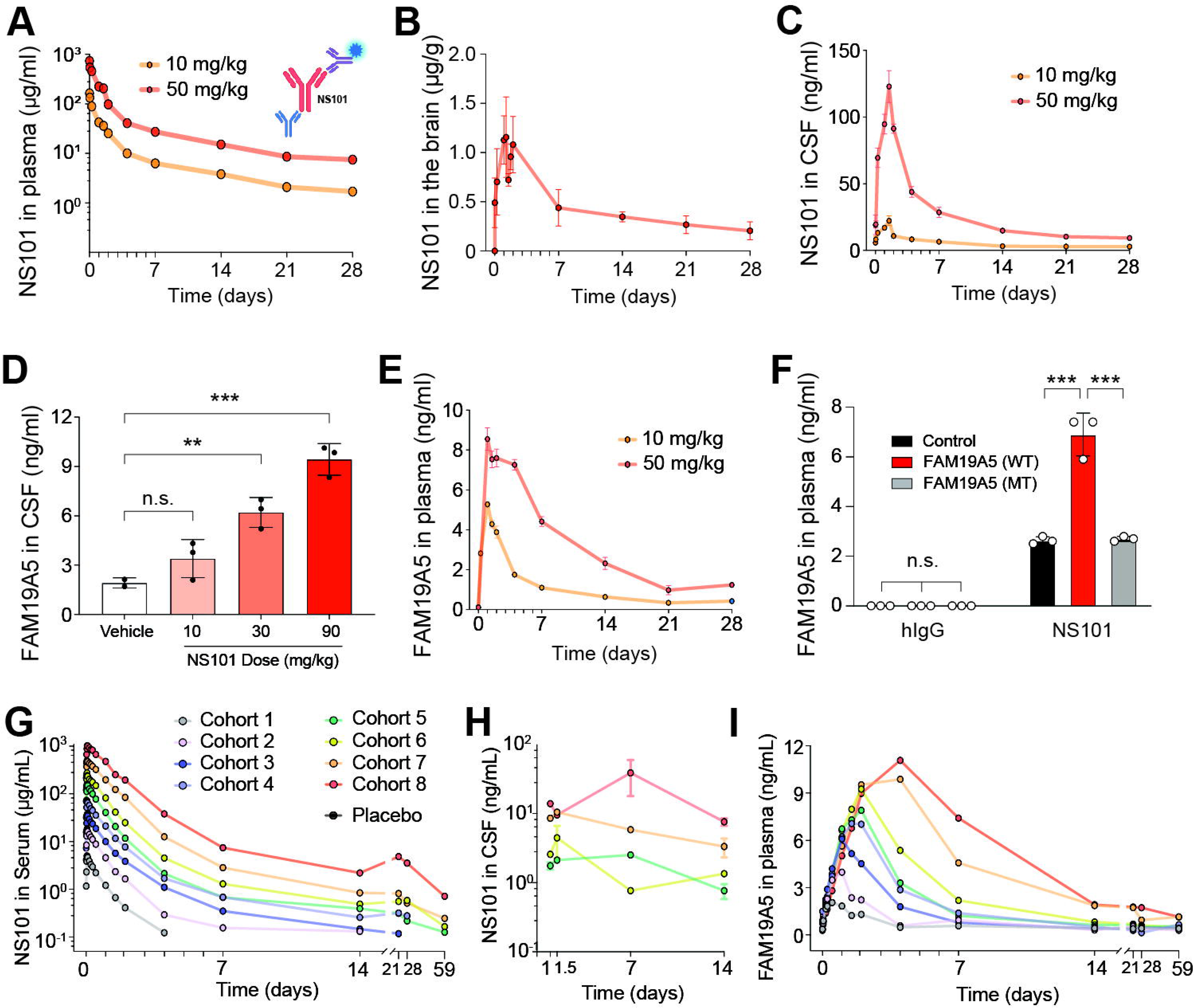
NS101, a potent monoclonal antibody against FAM19A5. (A) Plasma NS101 levels at the indicated time points after a single IV administration of 10 mg/kg or 50 mg/kg NS101 determined via ELISA (inserted, captured by rabbit anti-human IgG Fc, detected by HRP-conjugated hIgG kappa light chain antibody). NS101 levels in the (B) mouse brain and (C) rat CSF at the indicated time points after a single IV administration of 10 or 50 mg/kg determined via ELISA (captured by rabbit anti-human IgG heavy chain, detected by HRP-conjugated hIgG kappa light chain antibody). (D) Brain-secreted FAM19A5 levels in mouse CSF 48 hours after a single IV administration of 10, 30 or 90 mg/kg NS101 (captured by S-A5-Ab, detected by HRP-conjugated SS01). Data are presented as the mean ± SEM. One-way ANOVA followed by Tukey’s multiple comparison test was used to calculate *P* values. n.s., not significant; \*\**P* < 0.01, \*\*\**P* < 0.001. (E) Brain-secreted FAM19A5 levels in rat blood plasma at the indicated time points after a single IV administration of 10 or 50 mg/kg NS101, as determined via ELISA (inserted, captured by His-TEV hLRRC4B(453-576), detected by HRP-conjugated C-A5-Ab). (F) Plasma FAM19A5 levels after brain infusion of WT and MT FAM19A5 (R58A, R59A). One day after the infusion, the mice were IV administered hIgG or 10 mg/kg NS101. ELISA was performed using His-TEV LRRC4B and HRP-conjugated SS01 (n = 3 per group). Data are presented as the mean ± SEM. One-way ANOVA followed by Tukey’s multiple comparison test was used to calculate *P* values. n.s., not significant; \*\*\**P* < 0.001. (G-H) NS101 levels in (G) human serum and (H) human CSF at the indicated time points after a single IV administration of 0.25 to 48 mg/kg were determined via ELISA (captured by N’HIS-TEV FAM19A5 and rabbit anti-NS101, detected by sulfo-tag anti-rabbit antibody). (I) Brain-secreted FAM19A5 levels in human blood plasma at the indicated time points after administration determined by ELISA (captured by S-A5-Ab, detected by HRP-conjugated SS01).

To assess the target engagement of NS101, we measured CSF FAM19A5 levels 48 hours after a single IV injection of increasing doses of NS101. We observed a dose-dependent increase in the concentration of FAM19A5 in the CSF, indicating that NS101 bound to FAM19A5 in the brain and facilitated its release into the CSF (Figure 4D). Additionally, we tracked the concentration of FAM19A5 in plasma originating from the brain over 28 days. Typically, the concentration of FAM19A5 in plasma is negligible (Figure S7A). However, after a single administration of NS101, FAM19A5 levels in the plasma increased steeply for up to 2 days and then slowly decreased over 28 days in a dose-dependent manner (Figure 4E). These results suggested that FAM19A5 is released from the brain upon interaction with NS101, resulting in a plasma FAM19A5 profile that mirrors the concentration profile of NS101 in the brain and CSF. Additionally, infusing the brain with recombinant FAM19A5 amplified the levels of plasma FAM19A5 when NS101 was administered. In contrast, an infusion of MT FAM19A5 did not produce a similar effect (Figure 4F). This finding suggested that binding of FAM19A5 to NS101 facilitates its clearance from the brain, resulting in elevated plasma FAM19A5 levels and a consequent decrease in brain FAM19A5 levels. Notably, repeated administration of NS101 at high doses did not lead to any detectable adverse effects in terms of long-term toxicity (Figure S7B, C).

We then investigated the safety, pharmacokinetics, and target engagement of NS101 in a phase 1 clinical trial involving a total of 64 healthy subjects. The subjects were divided into 8 groups, with 2 subjects in each group receiving a placebo. The antibody dose was set at 0.25-48.0 mg/kg. The concentration of IV infused NS101 in the serum increased rapidly, with a median T_max_ of 0.98 to 1.74 hours across the doses. Subsequently, the NS101 levels slowly decreased over 60 days (Figure 4G). Antibodies delivered to the brain across the BBB were secreted into the CSF. NS101 secreted into the CSF was detected in the 6 mg/kg dose group and above, and the amount of NS101 detected tended to increase in proportion to the dose (Figure 4H). In the placebo group, only approximately 0.3 ng/ml plasma FAM19A5 was detected, and this level did not change noticeably over time. In contrast, the peak concentration and time to maximum concentration of plasma FAM19A5 significantly increased in a dose-dependent manner following the administration of NS101, as determined by the median time to maximum effect (median TE_max_ of 11.96 to 95.55 hours). At concentrations greater than 6 mg/kg, NS101 did not distinctly increase the peak concentration of FAM19A5 but increased the total amount of plasma FAM19A5 in a dose-dependent manner (the AUC ranged from 707843.52 to 7730521.14 hour·pg/mL) (Figure 4I). In addition, no dose-related adverse events, including serious side effects and death, were found in any of the experimental groups. These results demonstrate that NS101 can be delivered to the brain and target FAM19A5 in humans in the same way.

### 3.5 NS101 prevents synapse loss, increasing functional synapses in mouse models of AD

To investigate the potential association of FAM19A5 with AD pathology, we examined the levels of FAM19A5, LRRC4B, and PTPRF in P301S and APP/PS1 mice. Individual levels of these three proteins did not differ significantly between AD mice and their healthy littermate controls (Figure 5A and Figure S8). However, Co-IP experiments using an LRRC4B antibody revealed a significant increase in the amount of FAM19A5 bound to LRRC4B in both P301S and APP/PS1 mice compared to that in controls. Conversely, the level of PTPRF bound to LRRC4B decreased (Figure 5B). These findings suggest that in the AD brain, increased formation of the FAM19A5-LRRC4B complex may be associated with a reduction in synapses. Therefore, disrupting the FAM19A5-LRRC4B complex through NS101 treatment may offer a potential therapeutic strategy for restoring synapse number and function in AD.

**FIGURE 5.**
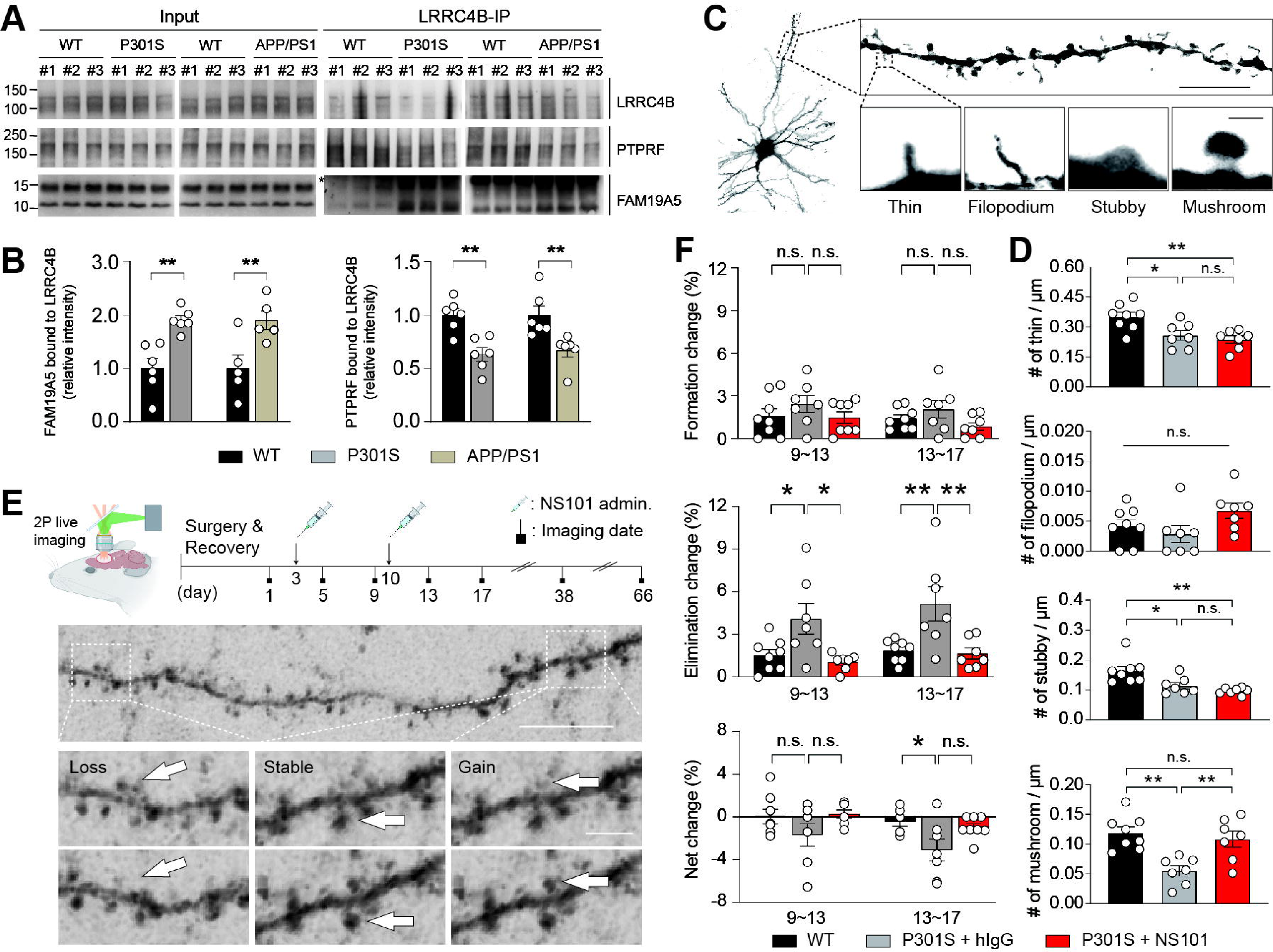
NS101 prevents synapse elimination, increasing the number of functional synapses in the AD model. (A) Co-IP using lysates prepared from the cerebral cortex of APP/PS1 mice, P301S mice, and their littermates to investigate the interaction between LRRC4B, FAM19A5, and PTPRF. (B) Relative band intensity of FAM19A5 bound to LRRC4B (left) and PTPRF bound to LRRC4B between the AD mice and littermates. Data are presented as the mean ± SEM. Unpaired t-test was used to calculate *P* values. **P < 0.01. (C) Representative Golgi staining image of a cortical neuron in the P301S mouse brain for spine classification. Scale bar, 10 μm (dendrite), 1 μm (spine). (D) Quantification of the number of different types of spines before and after NS101 treatment (n = 8 to 7 per group). Data are presented as the mean ± SEM. One-way ANOVA followed by Tukey’s multiple comparison test was used to calculate *P* values. n.s., not significant; \**P* < 0.05, \*\**P* < 0.01. (E) Representative image of spine dynamics in the P301S mouse brain. After cranial window surgery, dendritic spines in the somatosensory cortex were measured at the indicated time points. Scale bar, 10 μm (dendrite), 1 μm (spine). (F) Quantification of spine dynamics at days 9∼13 and 13∼17 in P301S mice. The formation and elimination rates and their net changes after the intraperitoneal administration of 30 mg/kg NS101 (n = 7 to 8 per group). Data are presented as the mean ± SEM. One-way ANOVA with Bonferroni’s multiple comparisons test was used to calculate *P* values. n.s., not significant; \**P* < 0.05, \*\**P* < 0.01.

We also measured the concentrations of FAM19A5 and tau proteins in CSF collected from humans of different ages. The concentration of FAM19A5 in human CSF increased with age (Figure S9A). Furthermore, the levels of CSF tau proteins, including phospho-tau (p-tau), the primary component of neuronal tangles in AD^43,44^, increased in parallel with the CSF FAM19A5 levels (Figure S9B). This finding aligns with the results of RNA data analysis in the brain tissue of AD patients^45^.

These results raises a hypothesis that the finely tuned synaptic balance, under the precise regulation of synaptolyitc factors including FAM19A5, disintegrates upon the influx of amyloid or tau aggregates, leading to excessive synapse loss. By inhibiting FAM19A5 with NS101 rather than aggregates, we aimed to rebalance the equilibrium, which could facilitate synapse and cognitive restoration. To investigate this possibility, 6-month-old P301S mice, a model of advanced AD exhibiting progressive synapse loss in the cortex due to the accumulation of p-tau proteins^46–48^, were IV administered 10 mg/kg NS101 weekly for 4 weeks. At 7 months of age, P301S mice showed a notable reduction in spine density in the prefrontal cortex compared to that of their WT littermates. In particular, there was a marked reduction in the number of mushroom spines, which are mature spines capable of signal transmission (Figure 5C, D). However, after NS101 administration, the mushroom spine density rebounded, reaching the level observed in WT mice (Figure 5D). Meanwhile, the densities of other spine types remained relatively unaffected by NS101 treatment. In contrast to the cortex, spine loss was not prominent in the hippocampal CA1 region of P301S mice at this age, as this region is relatively less affected by tauopathy in P301S mice than in WT mice^47,49^. Consequently, no spine restoration effects of NS101 were observed in this region (Figure S10).

Synaptic loss in P301S mice is driven by the toxicity of tau tangles, potentially leading to an increased rate of synapse elimination. Inhibiting FAM19A5 with NS101 may attenuate the elimination rate, rebalancing the disrupted synaptic balance, which likely contributes to the preservation of synapses and the subsequent increase in mushroom spines. To investigate whether there was a recovery in the balance of synapse formation and elimination after NS101 administration, we employed two-photon live imaging to trace spine changes in the somatosensory cortex of 7-month-old P301S mice following NS101 treatment for 66 days (Figure 5E). Notably, under physiological conditions, only a small fraction of spines are formed and eliminated at a similar rate^50^. Consequently, the net balance of spine formation and elimination reaches a steady state of zero, maintaining a stable state^51^.Compared to WT mice, P301S mice exhibited a significantly greater rate of spine elimination, while spine formation rates remained unchanged between the groups. Consequently, the net change increased significantly from 0% to 3.5%. Following two systemic injections of NS101 at weekly intervals, the spine elimination rate in P301S mice significantly decreased, resulting in the equilibrium of a net change comparable to that of WT mice (Figure 5F). The rebalancing effect on spine elimination persisted for up to 2 months but did not reach statistical significance (Figure S11).

### 3.6 NS101 restores synaptic activity and cognitive behavior in mouse models of AD

Next, we investigated whether the increase in mature spine density associated with the restoration of synaptic balance translated into enhanced synaptic functions in AD. APP/PS1 mice, which exhibit synapse loss associated with amyloid beta plaques^52,53^, were used to assess synaptic function after NS101 administration. Thirteen-month-old APP/PS1 mice were IV administered NS101 weekly for 4 weeks. Subsequently, we measured the miniature excitatory postsynaptic currents (mEPSCs) in the CA1 pyramidal neurons of the hippocampus. The frequency of mEPSCs, which arise from the spontaneous release of neurotransmitters, was significantly lower in APP/PS1 mice than in their WT littermates. However, NS101 treatment restored the mEPSC frequency to levels comparable to those of WT littermates (Figure 6A), consistent with the observations of increased intensities of synapse marker proteins in the hippocampus induced by NS101 (Figure S12). These findings, along with the slightly increased mEPSC amplitude (Figure 6B), suggest that NS101 strengthens synaptic function. Synaptic strengthening by NS101 was further confirmed through field excitatory postsynaptic potential (fEPSP) measurements, revealing enhanced fEPSP slopes and fiber volley amplitudes (Figure 6C). This facilitated the induction and maintenance of long-term potentiation (LTP) at normal levels in APP/PS1 mice (Figure 6D).

**FIGURE 6.**
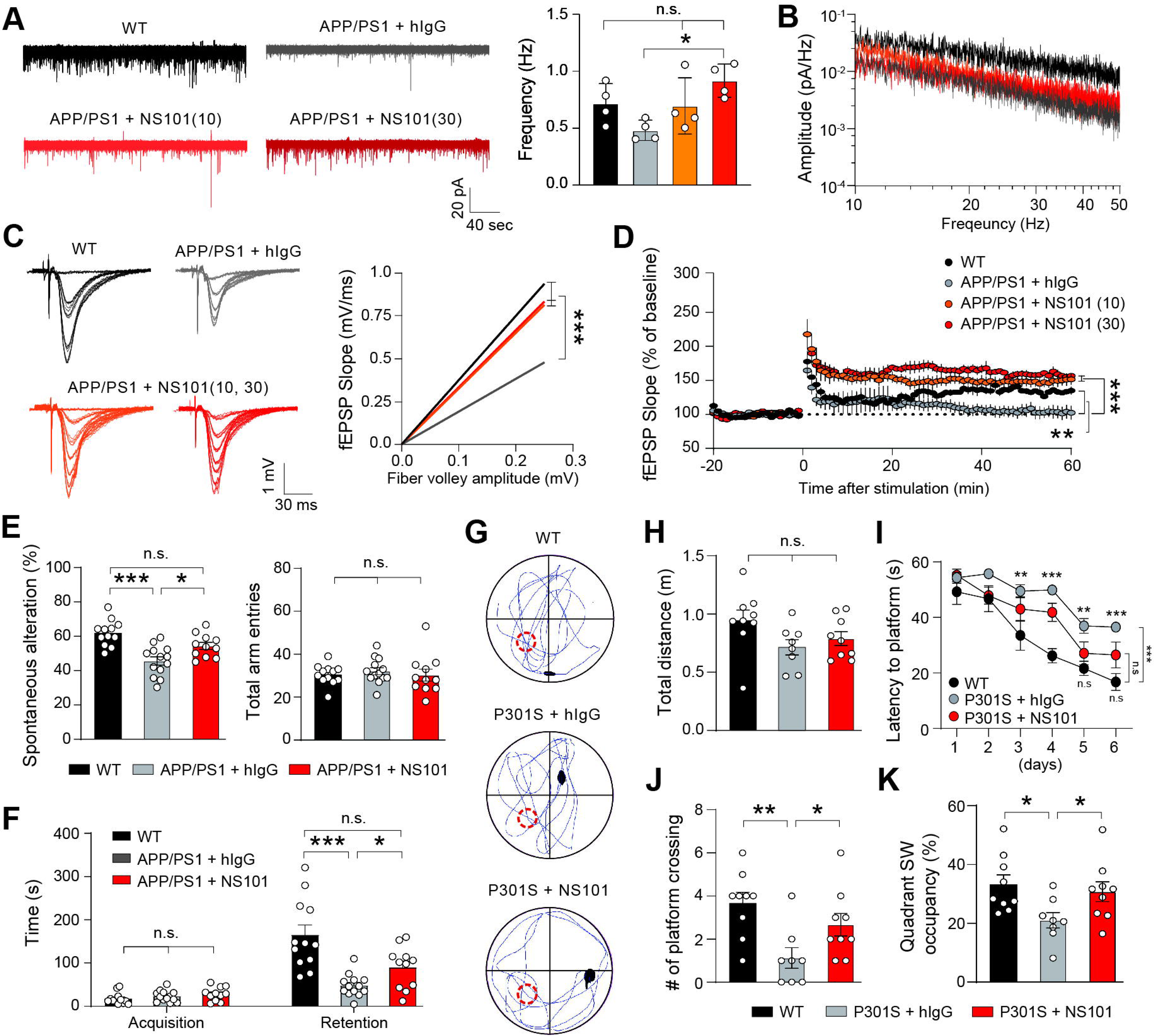
NS101 restores synaptic activity and cognitive behavior in AD. (A) Representative mEPSC traces in the hippocampal CA1 neurons of APP/PS1 mice and their frequency quantification (n = 4 per group). Data are presented as the mean ± SEM. One-way ANOVA followed by Tukey’s multiple comparison test was used to calculate *P* values. (B) Power spectrum analysis, displaying the average line of the dominating spikes before and after NS101 administration. (C) Representative input[output traces and fEPSP slopes plotted against fiber volley amplitudes in hippocampal CA1 neurons of WT and APP/PS1 mice treated with either hIgG or 10 mg/kg or 30 mg/kg NS101 (n = 5 per group). Data are presented as the mean ± SEM. One-way ANOVA followed by Tukey’s multiple comparison test was used to calculate *P* values. (D) The average fEPSP amplitude before and after theta burst stimulation to Schaffer collaterals for LTP induction (n = 5 to 6 per group). Data are presented as the mean ± SEM. One-way ANOVA followed by Tukey’s multiple comparison test was used to calculate *P* values. (E) Quantification of short-term memory in APP/PS1 mice using the Y-maze test (n = 11 to 13 per group). (F) Quantification of long-term memory in APP/PS1 mice using the passive avoidance task (n = 11 to 13 per group). Data are presented as the mean ± SEM. One-way ANOVA followed by Tukey’s multiple comparison test (for E and F) was used to calculate *P* values. n.s., not significant; \**P* < 0.05, \*\*\**P* < 0.001. Quantification of spatial learning and memory evaluated via the Morris water maze, (G) the swimming trajectories of WT and P301S mice, (H) the total distance traveled, (I) the latency to reach the platform, (J) the number of platform crossings, and (K) quadrant occupancy (n = 9 to 12 per group). Data are presented as the mean ± SEM. One-way ANOVA followed by Tukey’s multiple comparison test (for H, J, and K) and two-way ANOVA followed by the Newman[Keuls multiple comparisons test (for I) were used to calculate *P* values. n.s., not significant; \**P* < 0.05, \*\**P* < 0.01, \*\*\**P* < 0.001.

**FIGURE 7.**
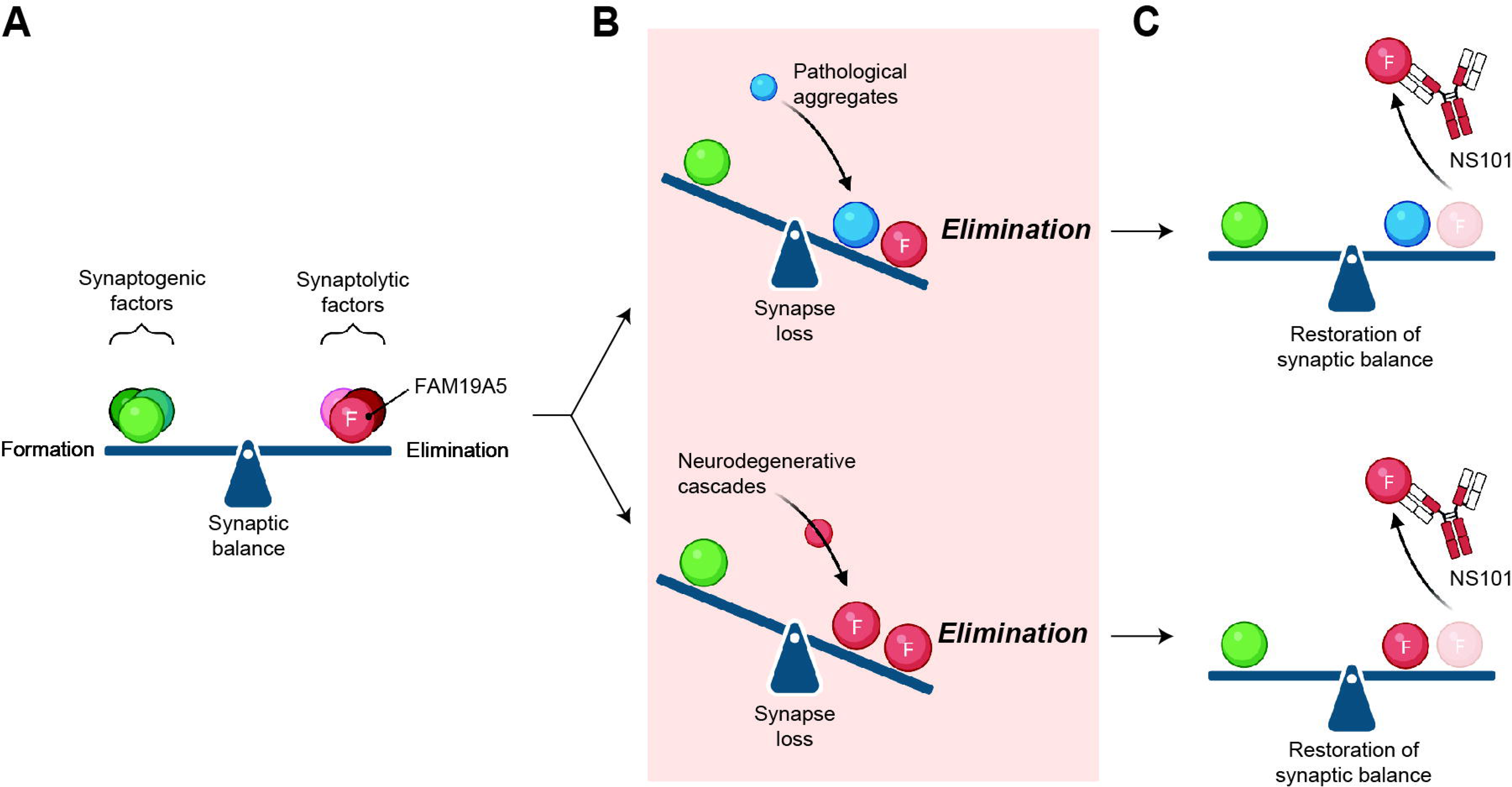
Illustration of synaptic balance restored by NS101. (A) Synaptic balance relies on the delicate interplay between synaptogenic and synaptolytic factors, ensuring the precise formation and elimination of synapses. (B) However, this equilibrium can be disrupted by the additional burden of aggregate toxicity or the upregulation of FAM19A5 by neurodegenerative cascades, leading to excessive synapse loss. (C) Targeting FAM19A5 with NS101 mitigates excessive elimination, restoring synaptic equilibrium and promoting synapse recovery.

We then investigated whether synaptic potentiation results in cognitive improvement. In the Y maze test, NS101 administration improved the spontaneous alternation rate of APP/PS1 mice to WT levels, while the total number of arm entries remained similar across all groups (Figure 6E). These findings suggested that NS101 restored short-term memory in the APP/PS1 mice. In the passive avoidance test, all groups showed no difference in initial memory acquisition to avoid electric shocks. However, NS101 administration significantly prevented the memory deficit observed in the APP/PS1 mice 24 hours after learning, resulting in memory levels that were similar to those of the WT mice (Figure 6F). These findings indicate that NS101 enhances the long-term memory of APP/PS1 mice. We then conducted the Morris water maze test with P301S mice (Figure 6G). All groups exhibited similar locomotion abilities, including swimming speed, as evidenced by the total distance traveled (Figure 6H). P301S mice took significantly longer to locate the platform than did WT mice. However, the NS101-treated P301S mice found the platform in a significantly shorter time (Figure 6I). The performance of the NS101-treated P301S mice was similar to that of the WT mice in terms of quantitative indicators such as platform crossing and spatial preference (Figure 6J, K). These results suggest that NS101 ameliorates the spatial learning and memory deficits associated with AD.

We then investigated whether the NS101-induced functional recovery of the AD brain is linked to the reduction of AD pathogens such as p-tau and Aβ. NS101 administration significantly reduced p-tau levels in the cerebral cortex but not in the hippocampus of P301S mice (Figure S13A, B). NS101 did not alter Aβ levels in either the cerebral cortex or hippocampus of APP/PS1 mice (Figure S13 C, D). These results in part align with the study by Suzuki et al.^36^, which demonstrated that a synaptic organizer protein itself can restore synaptic function and cognitive behaviors in an AD mouse model without affecting Aβ levels. We also measured microglial and astrocyte activation by immunostaining Iba1-and GFAP-positive areas in the brain. NS101 administration did not significantly alter microglial or astrocyte activation in either P301S or APP/PS1 mice, although there was a trend toward decreased Iba-1 and GFAP levels without statistical significance (Figure S13 E-L). Notably, the minimal effect on glial activation is consistent with the properties of NS101. To enhance its safety profile, NS101 was designed to minimize its affinity for Fc-gamma receptors on microglia, which are known for mAb-mediated toxicity^54^. Overall, these results suggest that NS101 exerted minor effects on the levels of AD pathogens and the activation of microglia and astrocytes but restored synaptic activity and cognitive function primarily by reestablishing synaptic balance through targeting FAM19A5.

## 4 DISCUSSION

Synapses undergo dynamic formation and elimination processes throughout life, maintaining a delicate balance that serves as a crucial mechanism for regulating neural plasticity^55,56^. However, in neurological diseases, the rate of synapse elimination exceeds the rate of synapse formation due to various pathological factors, resulting in a breakdown of synaptic balance and synaptic loss^22,57^. This synaptic loss generally occurs early in the disease process^58^, but even then, new synapses continue to form, albeit at a lower rate than the loss. This persistent formation suggests the potential for reversibility of synaptic balance and a critical window for intervention, offering hope for preventing or even reversing the disease process. Therefore, deepening our understanding of the mechanisms governing synapse dynamics may pave the way for more potent treatments for neurological diseases.

Synaptic adhesion molecules participate in regulating synapse numbers via molecular processes, including synapse formation, maturation, and continuous reorganization^55,59^. The role of synaptic adhesion molecules extends beyond just adhesive contacts between membranes, as they play a critical role in organizing synapses and modulating their functional integrity^60^. In particular, LRRC4B can form excitatory synapses via transsynaptic interactions with presynaptic protein tyrosine phosphatase receptors (PTPR-F, -D, and -S)^35,61,62^. Additionally, via its PDZ-binding domain, it binds to the postsynaptic scaffolding protein PSD95 and promotes clustering of NMDA and AMPAR receptors at the postsynaptic site^34,35^. Thus, LRRC4B plays a crucial role in various stages of excitatory synapse formation and function, such as inducing the initial transsynaptic connection between axons and dendrites, clustering postsynaptic proteins, and controlling synaptic plasticity^33,35^. Its widespread expression across various types of neurons in mice and humans has been revealed by recent single-cell RNA-seq^4,5^, further underscoring its role in general synaptic dynamics or balance.

This study identified FAM19A5 as a novel regulator of synaptic balance. Its strong binding affinity to the FB domain of LRRC4B disrupted the binding between LRRC4B and PTPRF, potentially triggering synapse disassembly. This hypothesis is supported by the observation that treating primary cultured neurons with FAM19A5 decreased the colocalization of synapse marker proteins. However, this effect was not observed for mutant FAM19A5, which lacks binding to LRRC4B. In addition, treatment with NS101, which can block the binding of FAM19A5 to LRRC4B, led to an increase in synapse number. These results suggest that, under physiological conditions, FAM19A5 plays a crucial role in maintaining the delicate equilibrium/balance between synapse formation and elimination, acting as a counterpoint to synaptogenic proteins (Figure 6A).

Synaptic balance can be disrupted in various neurological diseases^63^, including AD, where the Aβ and p-tau proteins and their harmful effects on neurons may contribute to significant synapse loss in the aging brain^22,64,65^. The lower spine number in tauopathy P301S mice compared to WT mice likely arises primarily from a significantly greater rate of spine elimination rather than a lower rate of spine formation. Treating P301S mice with NS101 to inhibit FAM19A5 normalized the spine elimination rate to that of WT mice, restoring the overall spine balance and leading to recovery of mature synapse numbers and function.

Notably, NS101 exhibited comparable efficacy in restoring synapse numbers within the hippocampi of APP/PS1 mice, a model of amyloidopathy. These findings suggest that inhibition of a single synaptolytic factor, FAM19A5, even in the presence of external synapse disrupting factors, such as amyloid or p-tau aggregates, can normalize the synaptic balance, restoring lost synapses and regaining impaired cognition across tauopathy and amyloidopathy conditions (Figure 6B, C).

FAM19A5 levels in the brain may increase as a result of neurodegenerative cascades, such as AD, aging, and brain injury^6–8,66^, representing another pathway leading to the loss of synapses (Figure 6B, C). Recent single-cell/nuclear RNA-seq analyses of human samples revealed increased FAM19A5 transcript levels in AD brain neurons^6^, especially in tau-tangle-bearing neurons^66^, indicating a potential link between FAM19A5 and AD. These findings are further supported by our current study, which demonstrates that FAM19A5 levels in human CSF increase with age in association with total and p-tau levels. Additionally, our earlier study showed increased transcript levels in a mouse model of traumatic brain injury^8^. Therefore, normalizing the synaptic balance by inhibiting the common synaptolytic factor FAM19A5 with NS101 offers a promising therapeutic approach for various neurological diseases characterized by excessive synapse loss arising from either external synaptolytic factors and/or increased FAM19A5 levels.

The recent success of a synthetic synapse organizer in directly restoring functional synapses and improving cognitive function in a mouse model of AD underscores the critical role of synapse restoration for AD treatment^36^. This suggests that even under adverse conditions, enhancing synaptic structure can normalize transmission, potentially triggering a positive feedback loop between healthy neurons and their environment. Notably, both this study and our own research offer structure-guided molecular tools as promising avenues for normalizing synapse function in patients with neurodegenerative diseases. An important advantage of our work over the synthetic synaptic organizer is the noninvasive delivery of NS101 into the brain. Systemic administration of NS101 crossed the blood[brain barrier and facilitated the transport of FAM19A5 from the brain to peripheral blood in mice, rats, and humans in a clinical trial. This consistency of effect across species is likely due to the 100% identical sequence of the FAM19A5 protein in these species.

The transport of target molecules from the brain to peripheral blood via an antibody-mediated approach was recently demonstrated in another study in which an anti-tau antibody captured brain tau proteins and transported them to the periphery in both transgenic mice and humans^67^. In this study, the transfer of target molecules from the brain to peripheral blood was readily apparent, as the baseline levels of the target molecules were much greater in the brain than in the periphery. These findings, along with our results, suggest that antibodies can be effective tools for removing target molecules in the brain, potentially offering a treatment approach for various neurological diseases. Given that a single IV administration of NS101 induces the transport of brain FAM19A5 to the blood over a month, antibody therapy has a clear advantage in clinical studies over other biologics that require direct administration to the brain and have a short half-life in the blood^36,68^.

In conclusion, our study unravels the critical role of FAM19A5 in regulating synaptic balance by binding to the FB domain of LRRC4B, promoting the elimination of synapses. This finding suggests the therapeutic potential for strategies that disrupt this interaction. Notably, both the anti-FAM19A5 antibody NS101 and the engineered protein/peptide containing the FB domain successfully blocked FAM19A5 binding to LRRC4B, highlighting promising avenues for restoring synaptic balance. These approaches hold particular relevance for neurological diseases suffering from progressive loss of synapses, which are mainly controlled by the FAM19A5-LRRC4B interaction.

## Acknowledgments

The authors acknowledge Eun Bee Cho, Yongwoo Jeong, and Sitaek Oh for assisting in manuscript preparation. The authors also acknowledge Richard Larouche and Hyun Jung Jung for organizing and conducting the clinical phase I trial. The authors also acknowledge Kyungho Seong for performing in silico residue scanning on the FAM19A5-FB complex using the Schrödinger Platform.

## Consent Statement

All participants provided written informed consent for the phase 1 trial and for the collection of CSF and blood samples.

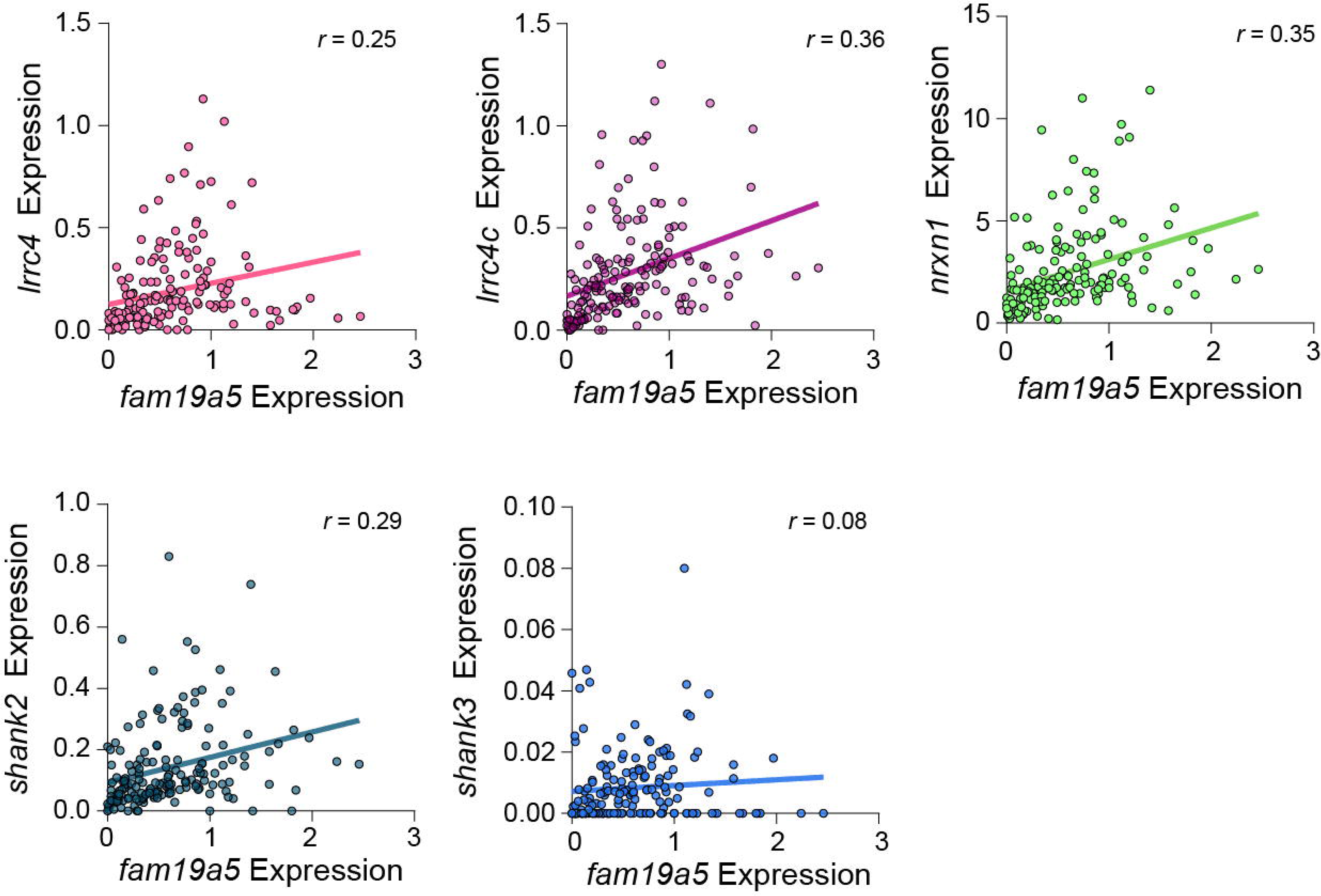

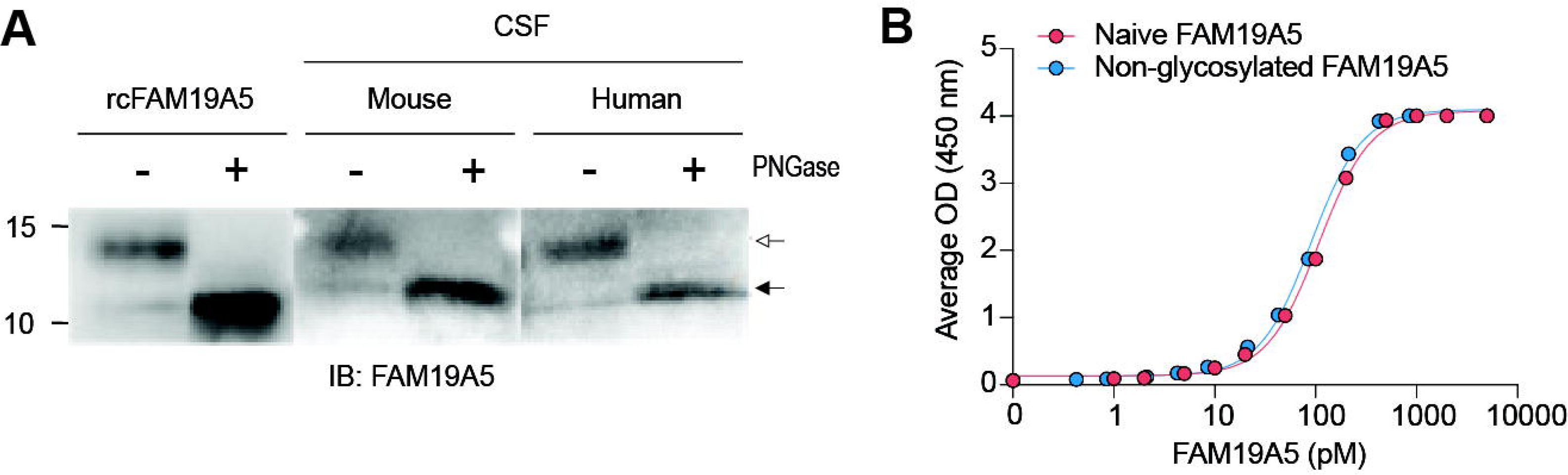

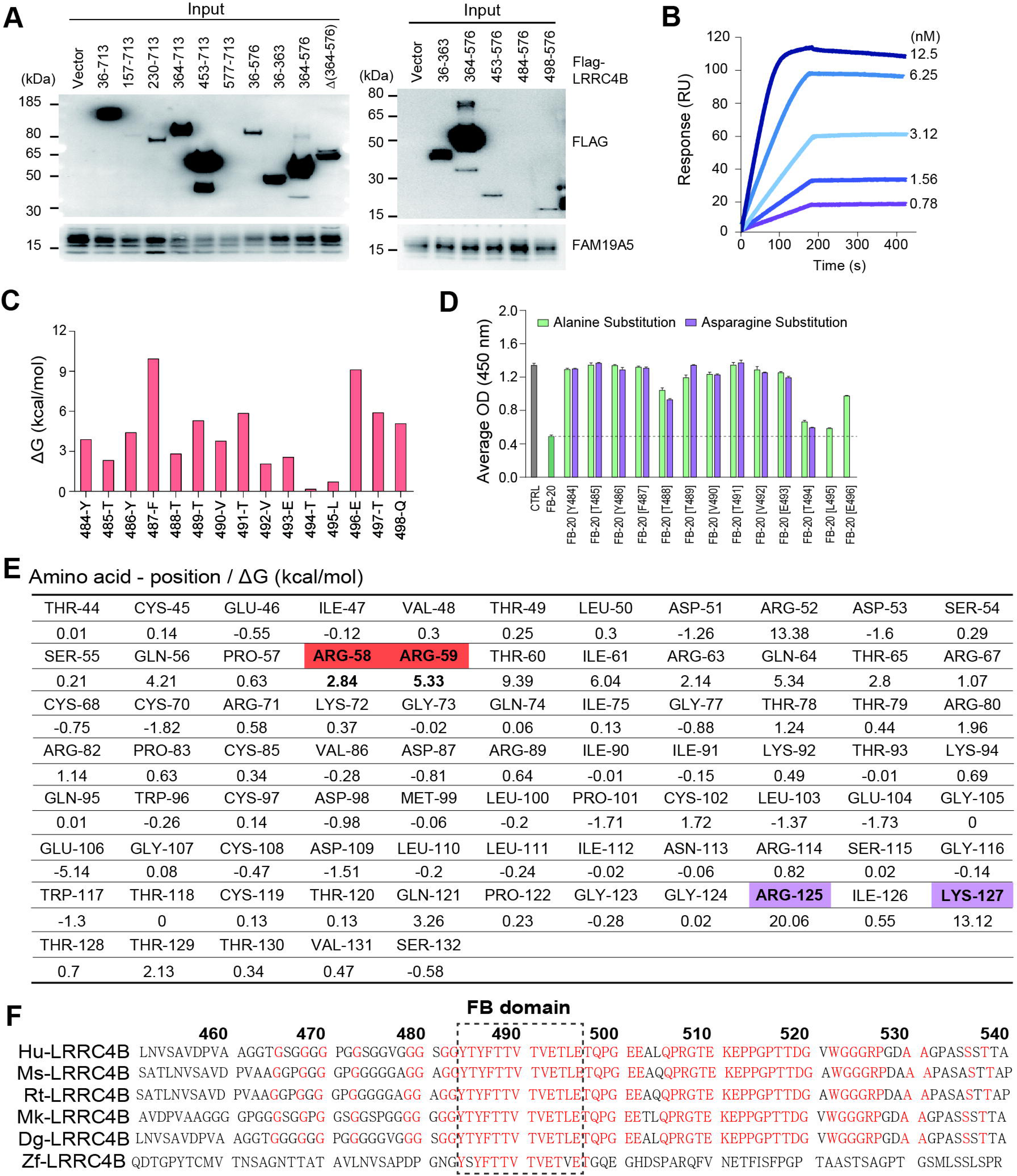

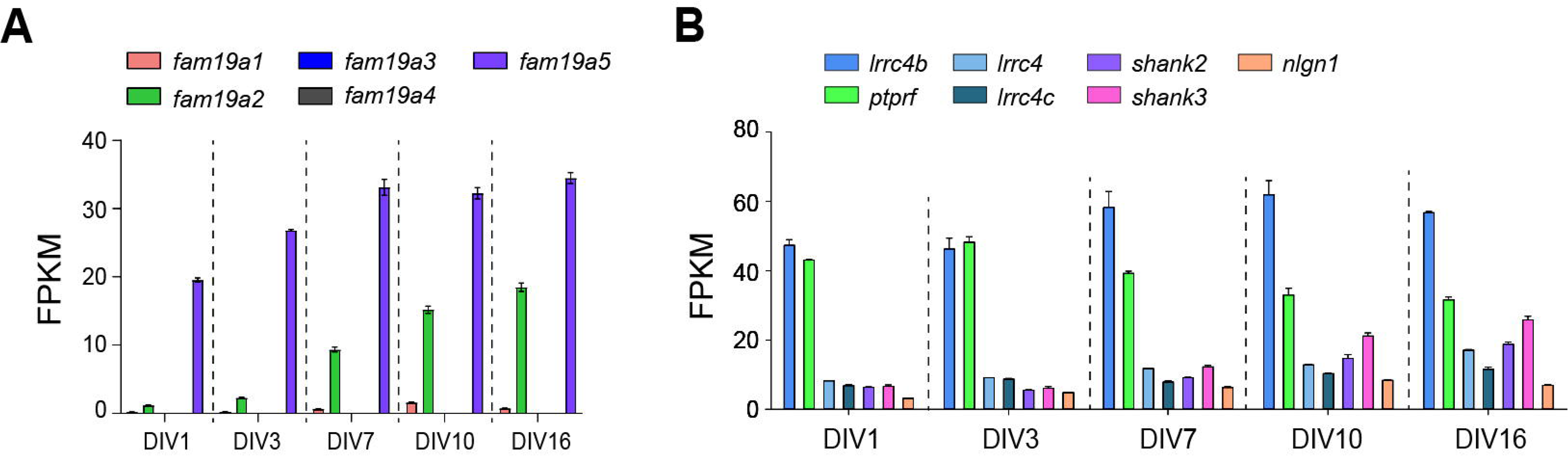

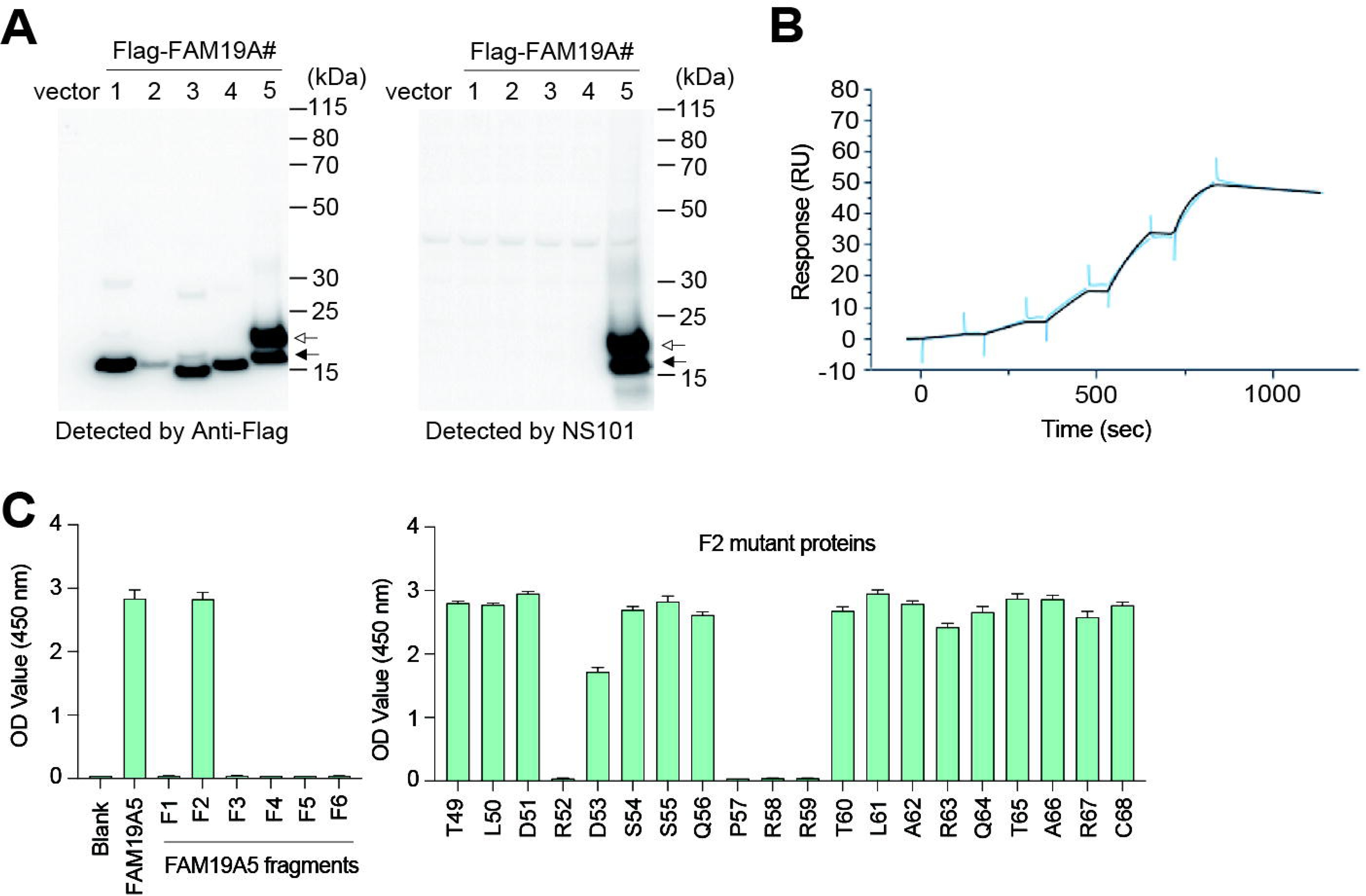

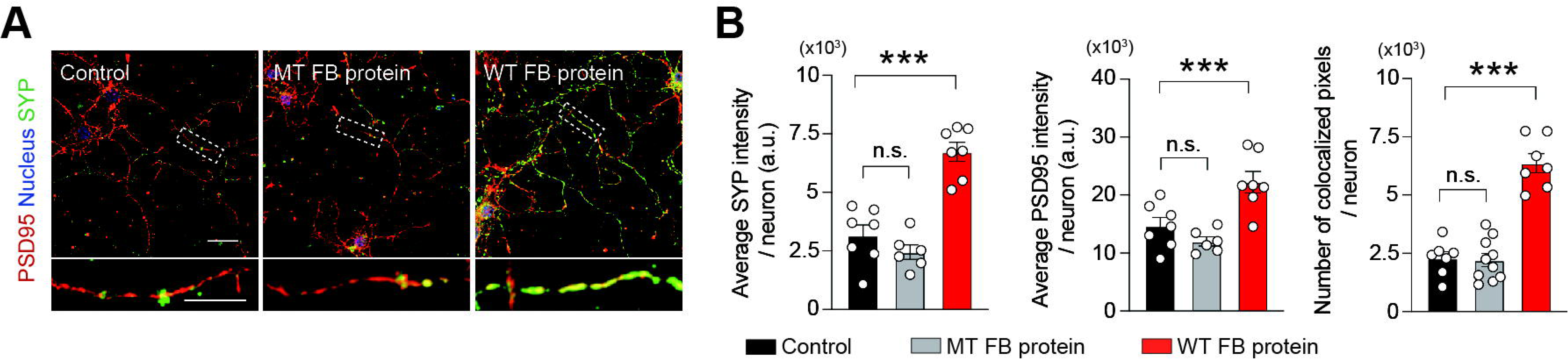

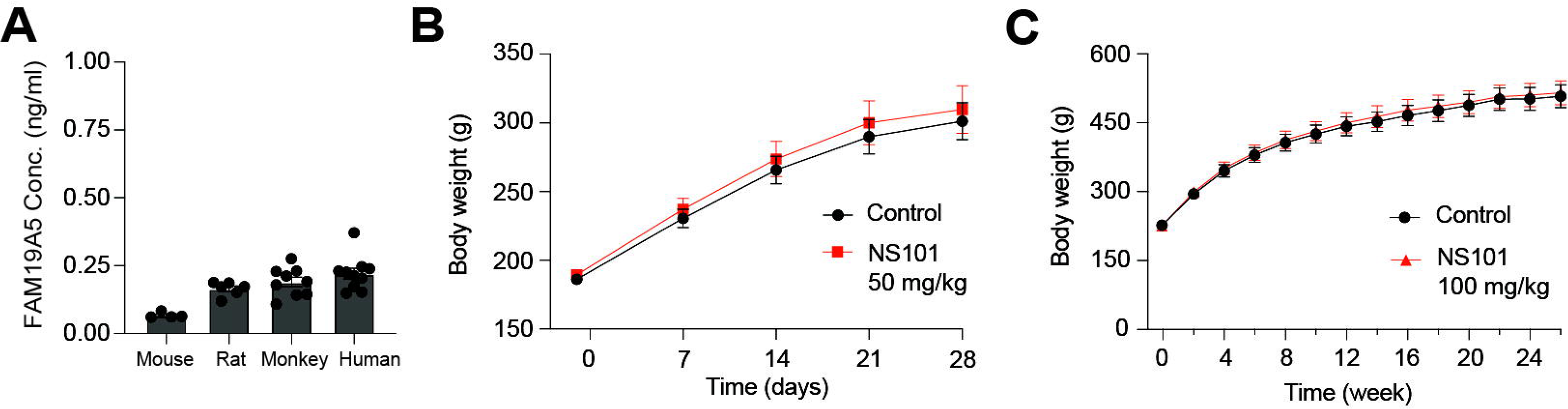

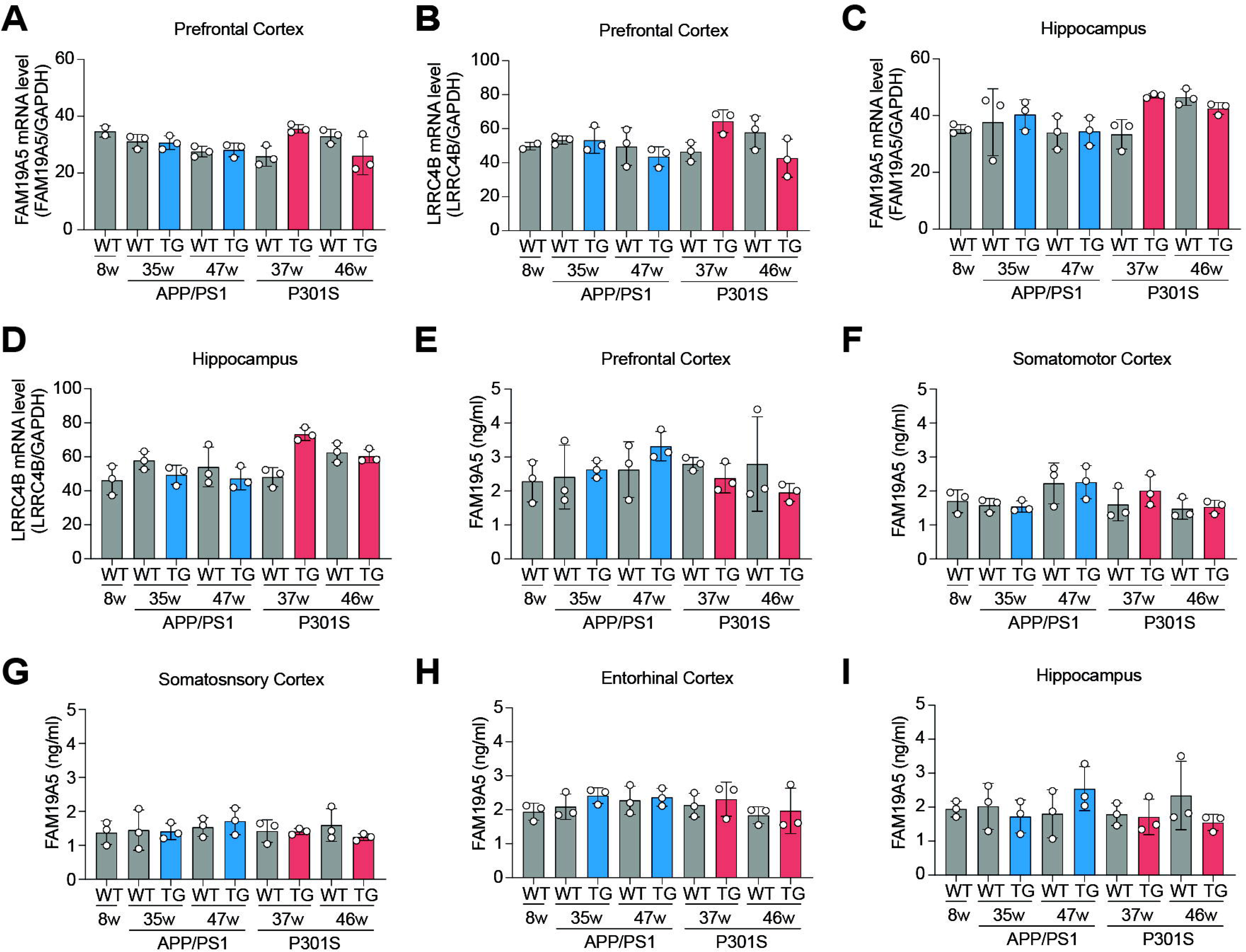

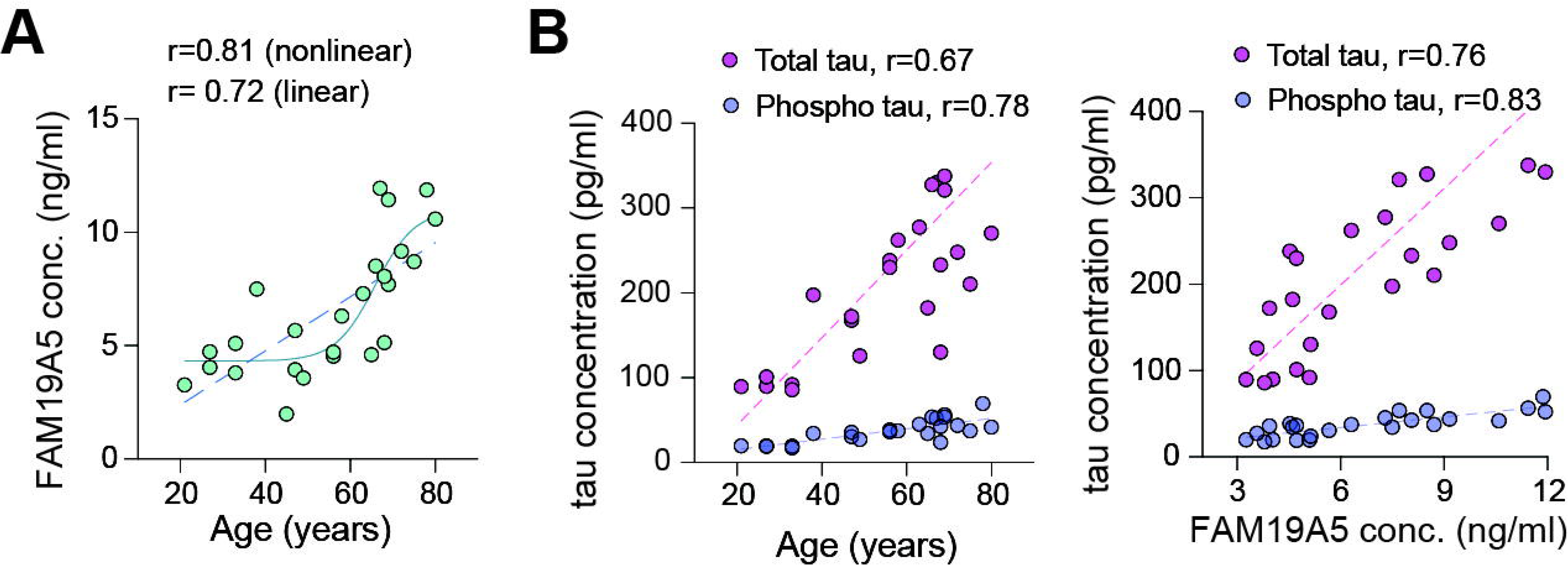

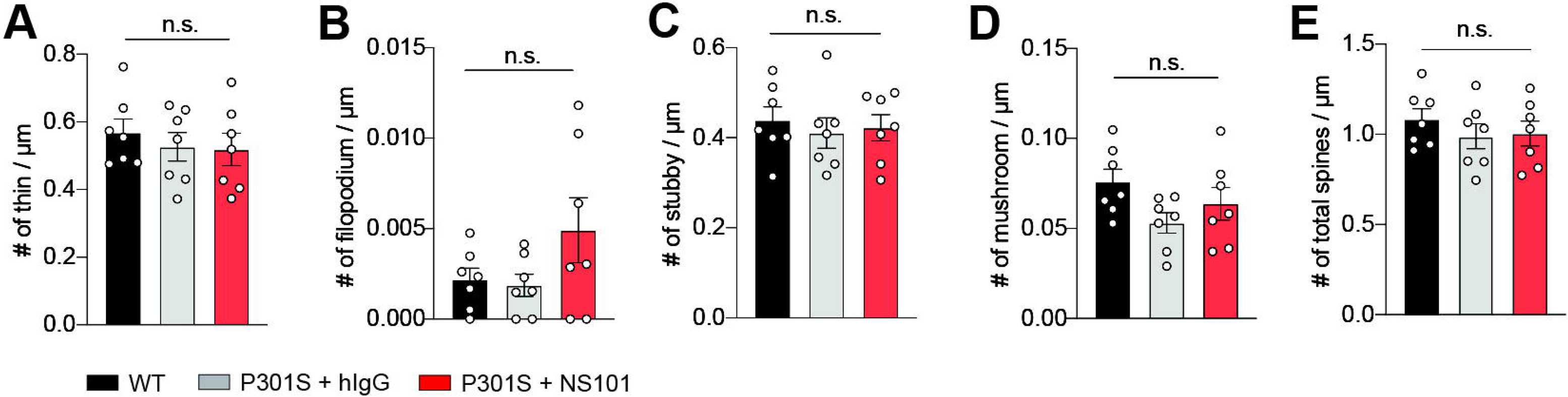

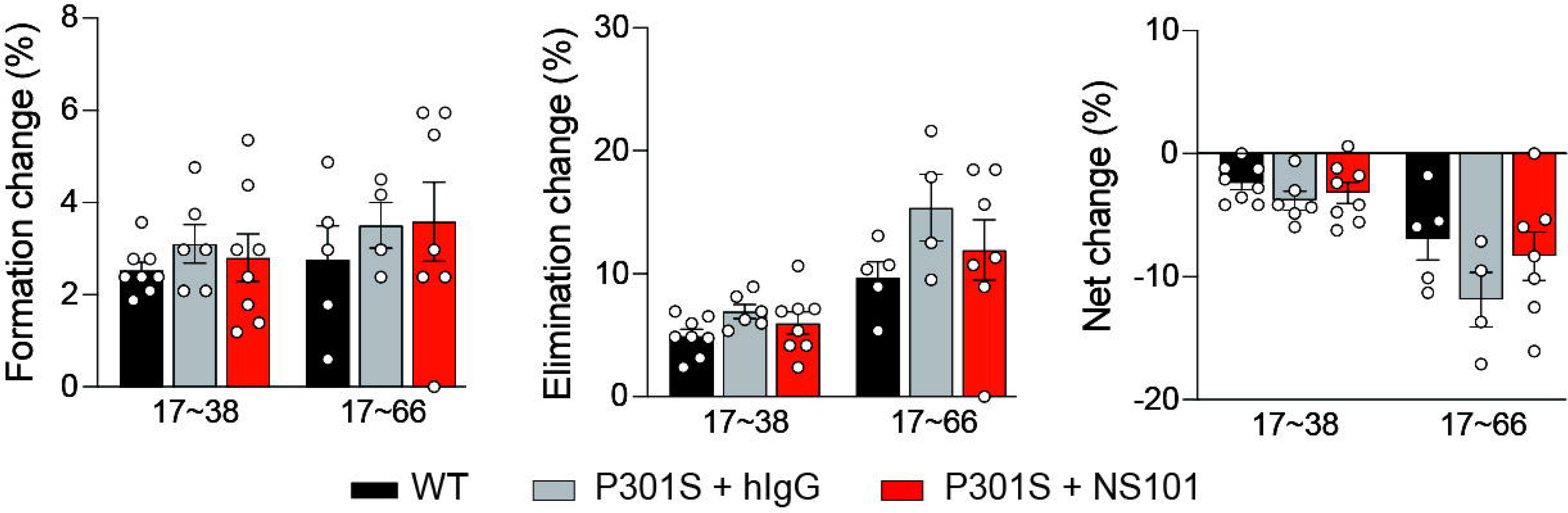

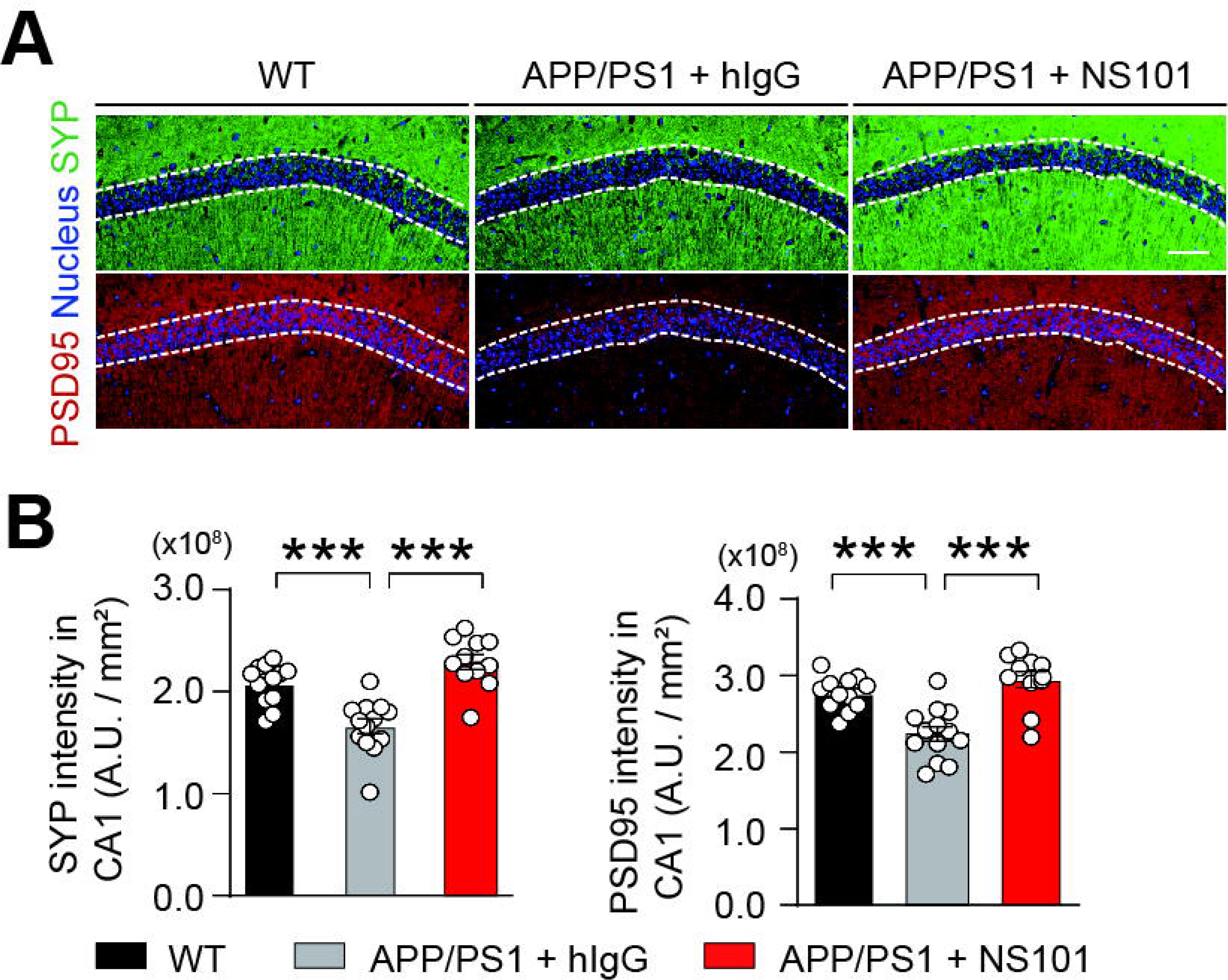

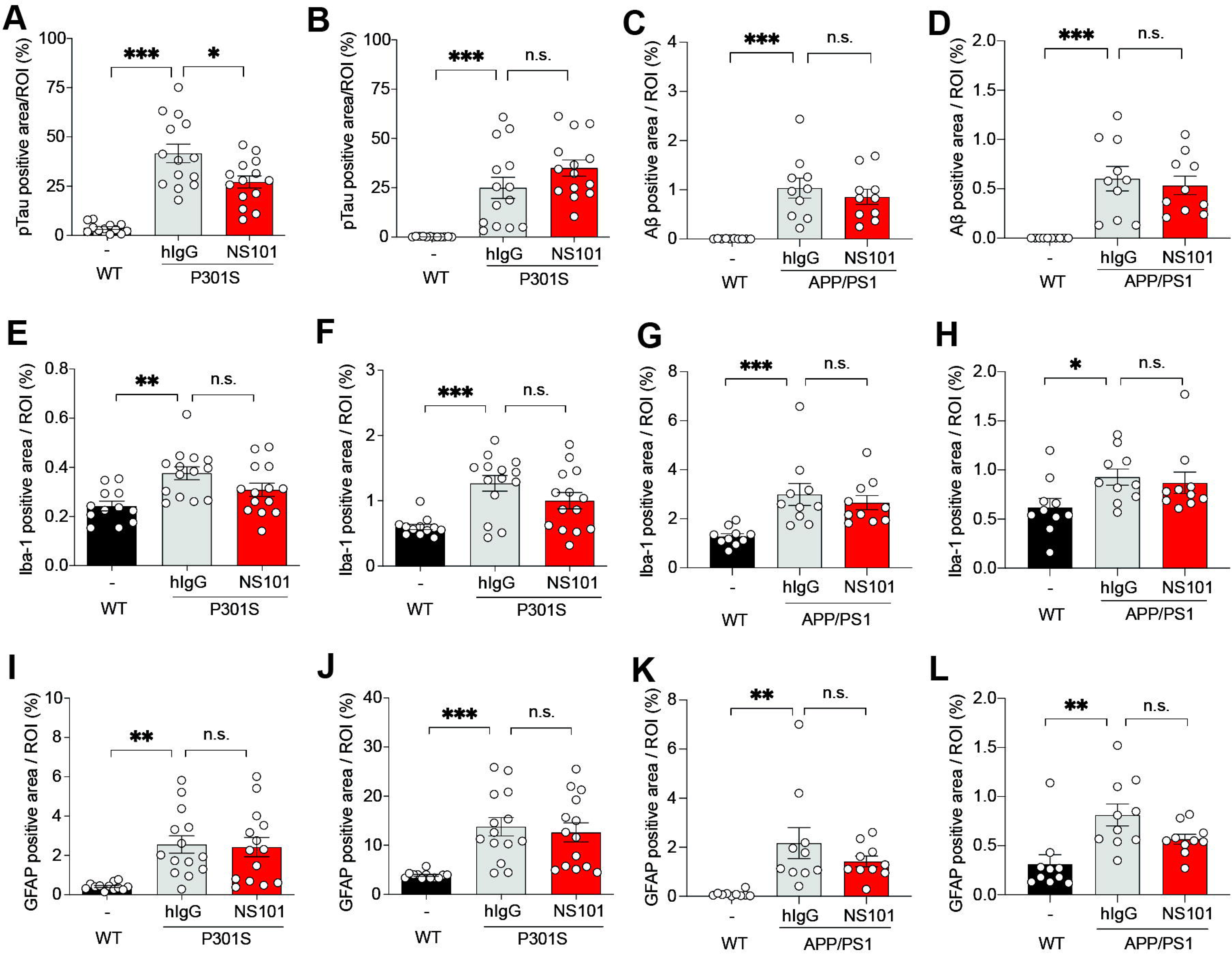

