## Supplementary Materials for "Inhibition of FAM19A5 restores synaptic loss and improves cognitive function in mouse models of Alzheimer’s disease"

**List of Supplementary Materials**

Extended Methods

Extended references

**Extended Methods**

Vertebrate animals

C57BL/6J mice (strain No. 000664), APP/PS1 [B6. Cg-Tg (APPswe, PSEN1dE9) 85Dbo/J] mice (Strain No. 034832-JAX), and P301S [B6. C3-Tg (Prnp-MAPT*P301S) PS19Vle/J] mice (strain No. 008169) were purchased from The Jackson Laboratory. Sprague‒Dawley rats were obtained from Orient Bio. The mice were housed under temperature-controlled conditions (22–23°C) with a 12-h light/12-h dark-light cycle (lights on at 8:00 am) and had ad libitum access to standard chow and water. The behavioral tests were conducted by the NDIC in accordance with the Animal Protection Act (Law No. 4372 enacted on May 31, 1991, partially revised Act No. 13023 on January 20, 2016) and approved by the Animal Experimentation Ethics Committee of KPC Co., Ltd. (Approval No. P191005). The procedures related to the analysis of animal samples were further approved by the Institutional Animal Care and Use Committee of Korea University (KOREA-2020-0031, KOREA-2020-0037, and KOREA-2021-0002). Ethical regulations were followed throughout.

Cell lines

HEK293 cells (ATCC# CRL-1573) were cultured in a humidified incubator at 37°C with 5% CO_2_. HEK293 cells were propagated in MEM with GlutaMAX^TM^ (Gibco) supplemented with 10% fetal bovine serum (Gibco) and 100 U/ml penicillin‒streptomycin (Gibco). For subculture, the cells were washed with DPBS (Gibco) and dissociated in 0.25% trypsin-EDTA (Gibco) for 5 min at 37°C.

DNA constructs

The pCMV20-Igκ_SP-FLAG vector was constructed by inserting a human Igκ signal peptide (SP) sequence in front of the FLAG tag of the pCMV20 plasmid (Sigma‒Aldrich). Full-length (36-713) and serial deletion constructs (36-363, 36-576, 157-713, 230-713, 364-713, 453-713, 577-713, 364-576, *364-576, 453-576, 484-576 and 498-576) of human LRRC4B (NM_001080457.1) were subcloned and inserted into the pCMV20-Igκ SP-FLAG vector. The pCMV20-Igκ SP-hFc vector contained Igκ SP at the N-terminus of the multiple cloning site (MCS) and human-Fc (hFc) at the C-terminus in the pCMV20 plasmid. The ectodomain (36-576) and fragment (453-576) of human LRRC4B were inserted into the MCS of the pCMV20-Igκ_SP-hFc vector. The mutant constructs in which Thr488 and Thr489 of LRRC4B were substituted with Ala or Ser were subcloned and inserted into the pCMV20-Igκ_SP-FLAG or pCMV20-Igκ_SP-hFc vector. FAM19A5 (WT and MT) and LRRC4B fragments (36-576 and 453-576) were inserted into the pCAG1.1-Igκ_SP-His-TEV vector. Mutant constructs in which Arg58, Arg59, and/or Arg125, Lys127 or Asn113 of FAM19A5 were replaced with Ala or Asp were subcloned and inserted into the pCAG1.1-Igκ_SP-His-TEV vector. Full-length human LRRC4B was subcloned and inserted into the pcDNA6-V5-His vector (Invitrogen). FAM19A5 (NM_001252310.1) without tags was subcloned and inserted into the pcDNA3.1 vector (GenScript). The ectodomain (30-1263) of PTPRF (NM_002840.4) was inserted into the pCMV20-Igκ_SP-hFc vector. All cloning was performed using the AccuRapid™ Cloning Kit (Bioneer) according to the manufacturer’s instructions. The primer sets used for cloning are listed in Table S1. Full-length human LRRC4B (OHu30422) and PTPRF (OHu02063) plasmid DNAs were purchased from GenScript.

Purification of recombinant proteins

Recombinant His-tagged proteins were purified according to methods described previously^1^. In brief, FAM19A5 (WT and MT) and LRRC4B (36-576) with N-terminal 6xHis tags, followed by a tobacco etch virus protease (TEV) recognition sequence, were cloned and inserted into the pCAG1.1 vector (a gift from Jong-Ik Hwang) and expressed in Expi293F cells (Gibco). The supernatant containing His-TEV-FAM19A5 was collected for Ni-NTA affinity chromatography (Cytiva). After washing with binding buffer containing 20 mM Tris-HCl (pH 7.5) and 200 mM NaCl, the proteins were eluted using binding buffers with increasing concentrations of imidazole (from 5 to 500 mM). Proteins eluted from 50 to 500 mM imidazole fractions were concentrated using Amicon® Ultra15 Centrifugal Filter Units (Millipore), and the buffer was exchanged for 1x DPBS (Gibco). The purified His-TEV-FAM19A5 proteins were digested overnight at 30°C with AcTEV protease (Invitrogen) to remove the His-tag and TEV sequence. The AcTEV protease containing the His-tag was then eliminated by overnight incubation with Ni-NTA, and digestion was confirmed using an anti-His tag antibody (Abcam). LRRC4B (ectodomain and fragments) and PTPRF (ectodomain) were cloned and inserted into the pCMV20-Igκ_SP-hFc vector and expressed in Expi 293F cells. The supernatant containing the hFc-fused proteins was collected using an rProtein A GraviTrap (Cytiva), and the hFc-fused proteins were eluted using an Ab buffer kit (Cytiva) following the manufacturer’s instructions.

Production of anti-FAM19A5 antibodies

We generated three chimeric chicken/human monoclonal antibodies against FAM19A5, named N-A5-Ab, C-A5-Ab, and S-A5-Ab, by immunizing chickens (Gallus gallus domesticus) with purified recombinant FAM19A5 using a previously described method^1^. N-A5-Ab and C-A5-Ab were further deimmunized and optimized by amino acid substitution to generate NS101 and SS01, respectively.

Production of protein fragments

To elucidate the key residues in the FB domain of LRRC4B involved in FAM19A5 binding, we generated custom-made mutant peptides in which each single residue was replaced with Ala or Asn. The amino acid sequences are listed in Table S2. To identify the epitopes of the FAM19A5 antibodies, the FAM19A5 linear sequence was divided into six groups, labeled F1 to F6. Further analysis was conducted to determine the crucial residue for binding, in which each single residue in F2 was replaced with Ala. The relevant proteins and their amino acid sequences are listed in Table S3 and Table S4.

Immunoblots

The cell lysates were prepared in lysis buffer containing 20 mM Tris-HCl (pH 7.5), 150 mM NaCl, 0.5% NP-40, and protease inhibitor cocktail (Thermo Scientific). The primary antibodies used are listed in Table S5. Immunoblots of the cell lysates were performed as described previously^1^.

For mouse brain tissue immunoblotting, mouse cerebral cortex tissues were homogenized in RIPA buffer (Thermo Scientific) supplemented with 25 mM Tris-HCl, 150 mM NaCl, 1% NP-40, 1% sodium deoxycholate, 0.1% SDS, and protease and phosphatase inhibitor cocktail (Thermo Scientific). The lysates were then boiled at 95°C for 5 min in SDS sample buffer (Biosesang) containing 100 mM DTT and subsequently loaded onto a 4–12% gradient Bis-Tris gel (Invitrogen) for electrophoresis. For all immunoblots, 20 μg of total protein were loaded per lane, fractionated by SDS-PAGE, and transferred onto PVDF (Thermo Scientific), which was blocked in 5% nonfat dried milk or 5% bovine serum albumin, followed by overnight incubation at 4°C with the appropriate antibodies: anti-FAM19A5 (N-A5-Ab), anti-LRRC4B (Alomone labs), anti-PSD95 (Invitrogen), and anti-PTPRF (Invitrogen). Signals were detected using SuperSignal West Pico PLUS Chemiluminescent Substrate (Thermo Scientific).

For the removal of N-glycosylation in FAM19A5, recombinant FAM19A5 (rcFAM19A5) and CSF samples were denatured and deglycosylated with the PNGase F enzyme according to the manufacturer’s instructions (New England Biolabs). In brief, 27 µl of CSF sample and 3 µl of 10X glycoprotein denaturing buffer containing 5% SDS and 400 mM DTT were combined and denatured by heating at 100°C for 10 minutes. Then, 4 µl of 10X GlycoBuffer 2 containing 500 mM sodium phosphate at pH 7.5, 4 µl of 10% NP-40 and 2 µl of PNGase F were added and incubated at 37°C for 1 hr. For immunoblotting, naïve samples and PNGase F-treated samples were resolved on Tris-glycine gels (Invitrogen).

Quantitative real-time polymerase chain reaction (qRT‒PCR) analysis

Total RNA from the mouse cerebral cortex and hippocampus was extracted with TRI Reagent (Invitrogen). Complementary DNA (cDNA) was obtained using a High-Capacity cDNA Reverse Transcription Kit (Applied Biosystems). The primer sequences are listed in Table S6. The CFX96 Touch RT‒PCR detection system using SsoAdvanced Universal SYBR Green Supermix (Bio-Rad) was used for qRT‒PCR. FAM19A5 and LRRC4B expression levels were normalized to the GAPDH level, and the relative quantity of mRNA was calculated using the comparative Cq method.

Phase 1 trial of NS101

The study NCT05143463 is a phase 1, double-blind, randomized, placebo-controlled, first-in-human (FIH) clinical trial designed to evaluate the safety and tolerability, pharmacokinetic (PK), pharmacodynamic (PD), and immunogenicity profiles of the study NS101 in healthy subjects under fasting conditions. We selected a healthy male volunteer population for this study because healthy subjects without any concomitant diseases or medications are a homogenous population, which allows proper evaluation of the safety, tolerability, and PK of NS101 without confounding factors. We assessed the subjects against the inclusion and exclusion criteria, which were described in our clinical protocol, to determine whether they were eligible to participate in this study. All subjects were provided with informed consent forms (ICFs) in their preferred language (either French or English) for review. Prior to initiation of the study procedures, the ICF was verbally reviewed with the subjects by qualified staff, allowing sufficient time for review of the information provided and to answer any questions the subjects had. Subjects were informed of any developments or changes to procedures that could influence their continued participation in the study. Subjects did not anticipate any direct benefits from participation in this research study, with the exception that they received a health evaluation. Subjects who participated in this study were compensated for their time; however, they were not offered any incentives.

NS101 infusion doses of 0.25, 0.75, 1.5, 3.0, 6.0, 12.0, 24.0, and 48.0 mg/kg (cohorts 1 to 8) were selected based on nonclinical pharmacology, PK, and toxicology studies in rats and monkeys. The study consisted of 8 cohorts, each including 8 subjects, randomized at a 3:1 ratio (6 subjects received NS101, and 2 received matching placebo), for a total of 64 subjects. For every cohort, the body weight of each subject measured the day before was used to calculate the exact individual dose required on a mg/kg basis based on their assigned dose level. The drug was administered on the morning of the day after the subjects had fasted overnight for at least 8 hours. The cohorts were dosed sequentially in an ascending fashion. The IV infusion was performed over a period of approximately 60 minutes in bed using aseptic techniques at a constant rate using a volume-controlled infusion device. At the end of the infusion, 3 mL of saline solution was injected to flush the remaining drug into the IV catheter. The end of the infusion was set to the end of the 3 ml flush.

In each cohort, a total of 21 blood samples were drawn into blood collection tubes (1 × 3 mL) containing plastic serum spray-coated silica before infusion and at 0.25 (±3 min), 0.5 (±3 min), 0.75 (±3 min), 1 (±3 min), 1.25 (±3 min), 1.5 (±3 min), 2 (±3 min), 4 (±3 min), 6 (±3 min), 8 (±3 min), 12 (±15 min), 24 (±15 min) (Day 2), 36 (±15 min) (Day 2), 48 (±15 min) (Day 3), 96 (Day 5±1), 168 (Day 8±1), 336 (Day 15±1), 504 (Day 22±2), 672 (Day 29±2), and 1416 (Day 60±3) hours after the start of infusion. Blood samples were kept at room temperature and centrifuged at 1300 ±20 g for at least 10 minutes at room temperature (no more than 180 minutes passed between the time of each blood draw and the start of centrifugation). Two aliquots of at least 0.5 mL (when possible) of serum were dispensed into polypropylene tubes as soon as possible. The aliquots were stored at -80°C until ELISA.

Subjects assigned to Cohorts 5 through 8 were randomized for collection of a CSF sample at one of the following timepoints: 24 (±3 horus), 36 (±3 horus), 168 (day 8±1), or 336 (day 15±1) hours poststart of infusion. Each of the 4 timepoints was assigned to 2 subjects. The sentinel subjects were randomized to the same timepoint to ensure that each timepoint was assigned to at least 1 subject receiving NS101. A single CSF sample was collected via lumbar puncture. The volume of collected CSF per sample did not exceed approximately 6 ml. A stabilizing agent (10% BSA in 1X PBS with 5% Tween-20) was added and mixed with the NS101 CSF samples (no more than 8 minutes between sample collection and mixing with the stabilizing agent). Two aliquots of at least 0.5 mL (when possible) of CSF were dispensed into polypropylene cryovials as soon as possible. The aliquots were stored at -80°C (±10°C) until ELISA.

The clinical study protocol, any relevant associated documents, and ICFs were reviewed and approved by an Institutional Review Board (IRB, **#**00000971) prior to beginning the associated study procedures. The ethics committee that reviewed the study was Advarra, an independent service provider located in Aurora, Ontario, Canada. The IRB had no representatives from Syneos Health or Sponsor and was, therefore, completely independent. All clinical work was conducted in compliance with Good Clinical Practices (GCPs) as referenced in the International Council for Harmonization (ICH) guidelines (ICH E6), Good Laboratory Practices (GLPs) as referenced in the ICH guidelines, and all applicable regulations, including the Federal Food, Drug and Cosmetic Act, U.S. applicable Code of Federal Regulations (CFR) Title 21, and any IEC requirements related to clinical studies.

Surface plasmon resonance (SPR)

SPR experiments were carried out on a Biacore 8K (Cytiva, for affinity between NS101 and FAM19A5) or a Biacore t200 (Cytiva, for affinity between FAM19A5 and LRRC4B) with active temperature control at 25°C following the manufacturer’s protocols. The running buffer was 1 × HBS-EP (10 mM HEPES, 150 mM NaCl, 3 mM EDTA, 0.05% Tween 20, pH 7.4). For immobilization, 6xHis-LRRC4B (453–576) in 10 mM sodium acetate, pH 4.5, which is a buffer equivalent to 0632–638 resonance units (RUs), was injected onto a nitrilotriacetic acid (NTA) chip at a 30 µl/min flow rate. FAM19A5 at increasing concentrations (0.78, 1.56, 3.12, 6.25, and 12.5 nM) was diluted in running buffer and flowed across the immobilized 6xHis-LRRC4B (453–576) for 180 s at a flow rate of 30 μl/min (association). The sample was replaced with running buffer for 240 s (disassociation). The chip surface was regenerated with 350 mM EDTA and 500 mM HBS-P imidazole. For all the samples, the blank injection with buffer alone was subtracted from the resulting reaction surface data. The data were analyzed using Biacore 8K evaluation software (Cytiva).

In silico modeling of FAM19A5-LRRC4B

We used AlphaFold2 to model the structure of the FAM19A5-LRRC4B complex. Input paired multiple sequence alignments (paired MSAs) were generated following the input generation protocol of RoseTTAFold. hMSAs for the FAM19A5 protein and LRRC4B extracellular domain were generated by an iterative sequence search against the UniClust30 database^2^ using HHblits^3^. To predict complex structures, we generated paired MSAs based on individual MSAs by pairing sequences from the same species. To provide pseudomultimer inputs to AlphaFold2, we used a gap insertion technique that has been previously applied to RoseTTAFold. The final models were ranked by the predicted TM score (pTM score), and the best scored model was used for further study after structural relaxation using Rosetta. To identify the salt bridge, PyMOL (Schrödinger, LLC) was used to identify the charged residues in the FAM19A5-LRRC4B complex. Then, atomic bonds between oxygen and nitrogen within 2.5-4.0 Å were screened.

In silico residue scanning of the FAM19A5-FB complex

We used the residue scanning module (Schrodinger Bioluminate®) for in silico residue scanning (e.g., Ala scanning) and to calculate the perturbation of protein‒protein binding affinity, which was defined as the change in binding free energy. To evaluate the potential impact of a mutation, we used ΔAffinity and ΔStability (solvated) values.

Primary neuronal culture

Primary cultured neurons were prepared from postnatal C57BL/6 pups (Nara Biotech) on postnatal day 1 following established procedures^4^. Initially, the cortices were dissected in Hank’s buffered salt solution (HBSS) (Invitrogen) and then subjected to digestion with 2.5% trypsin for 15 minutes at 37°C. After this step, the supernatant was removed, and the tissues were washed with HBSS. The tissues were gently triturated, and the dissociated cells were plated on glass coverslips precoated with poly-D-lysine (in borate buffer at a concentration of 50 μg/ml; Sigma‒Aldrich). The plated cells were housed in 60 mm culture dishes and cultured in minimum Eagle’s medium (MEM) supplemented with 0.5% glucose, 1 mM pyruvate, 1.2 mM L-glutamine, and 12% fetal bovine serum. After 6 hours of incubation, the medium was replaced with neurobasal medium (Invitrogen, Carlsbad, CA, USA) supplemented with 2% B-27 and 0.5 mM L-glutamine (Gibco). The cells were then maintained in a 5% CO_2_ humidified incubator at 37°C. Every 3–4 days, half of the medium in the culture dish was replaced with fresh culture medium.

For RNA-seq analysis, neurons were cultured for 16 days, and total RNA was isolated at various time points: 1, 3, 7, 10, and 16 days. The RNA was then subjected to RNA-seq analysis following established protocols^5^.

For the transfection of hippocampal neurons, a calcium phosphate transfection kit (Invitrogen) was used according to the manufacturer's instructions. Briefly, 10 µg of DNA was mixed with 2 M CaCl_2_ and diluted with HEPES-buffered saline (HBS), followed by incubation at room temperature for 30 minutes. The mixture was then applied to the prepared hippocampal neurons by dropwise addition. After transfection, the cells were fixed with 4% formaldehyde for immunocytochemistry (ICC).

Immunocytochemistry and immunohistochemistry

Immunocytochemistry for Primary Cultured Neurons: To conduct immunocytochemistry on primary hippocampal neuron, fixation was carried out with 4% PFA. After three subsequent washes with ice-cold DPBS, the cells were blocked using buffer containing 3% bovine serum albumin (BSA) and 0.1% Triton X-100 in PBS for 30 minutes. The cells were then incubated with primary antibodies overnight at 4°C. After three additional washes with DPBS, the cells were incubated with secondary antibodies and 20 mM Hoechst solution (Invitrogen) for 1 hour at room temperature. Following three final DPBS washes, we captured fluorescence images using a confocal microscope (Leica), and the coverslips were mounted with mounting solution (Biomeda). All the images were processed using Las X software (Leica). To quantify functional synapse, hippocampal neurons were transfected with EGFP in 10 DIV as described in above. At 13 DIV, neurons were fixed and immunostained with anti-SYP (Sigma‒Aldrich), multi-point tool and simple neurite tracer plugin, ImageJ (NIH) was used for quantification. To quantify pre- and postsynaptic molecules, cultured neurons were treated with wild type or mutant FB proteins, at DIV 3 and DIV 6. At DIV 7, the neurons were fixed and immunostained with anti-SYP (Sigma‒Aldrich) and anti-PSD95 (Invitrogen). The quantification of synaptic markers and their colocalization was performed using ImageJ (NIH) with the colocalization plugin.

Immunohistochemistry of brain slices: In the context of immunohistochemistry, mice underwent transcardial perfusion with a solution of 4% PFA in PBS, and their isolated brains were subsequently postfixed in the same fixative for 24 hours. These brains were then cryoprotected in 30% sucrose, serially sectioned on a cryostat (40 mm), and stored in a mixture of 50% glycerol and 50% PBS at −20°C until further use. The brain sections were blocked for 30 minutes in buffer composed of 3% BSA and 0.1% Triton X-100 in PBS. The membranes were then incubated with primary antibodies diluted in blocking buffer overnight at 4°C. The primary antibodies used included anti-SYP (Sigma‒Aldrich), anti-PSD95 (Invitrogen), anti-Tau (p-Ser396, Genetex), anti-β-amyloid (1-16, BioLegend), anti-Iba-1 (Wako), and anti-GFAP (Invitrogen) antibodies. After this overnight incubation, the sections were subjected to three PBS washes and then incubated with secondary antibodies or Hoechst 33242 diluted in PBS for 30 minutes at room temperature. Subsequently, the sections were washed again, mounted, and observed using either a slide scanner (Zeiss) or a confocal microscope (Leica). The quantification of immunohistochemistry images was performed using Las X (Leica) or ZEISS ZEN 2.6 (Zeiss) software. For the analysis of synaptic proteins (synaptophysin and PSD95), the mean intensity (A.U./mm²) was calculated by measuring the total intensity and dividing it by the sum of processed pixels in mm². For the analysis of pTau, Aβ, Iba-1, and GFAP, the relative area (%) of the positive signal was calculated by measuring the area of the positive signal and dividing it by the total region of interest (ROI) area.

Golgi staining and spine classification

Golgi staining was performed according to the manufacturer's instructions (FD NeuroTechnologies, PK401). Whole fresh brains were collected and placed in impregnation solution. The solution was replaced after 24 h, and then the brains were incubated at room temperature in the dark for 2 weeks with gentle shaking every 3 days to prevent precipitation of the impregnation chemicals. After the impregnation step, the brains were placed in a rehydration solution for 1 week. The brains were frozen in a container with dry ice-cooled isopentane (Sigma‒Aldrich, M32631) and cryosectioned into 100 μm slices. Prefrontal cortex slices were mounted on gelatin-coated slides (FD NeuroTechnolgies, PO101). For the developing step, the slides were incubated for 10 min in staining solution. The slides were dehydrated with increasing concentrations of pure ethanol (Sigma-Aldrich, E7023), cleared with xylene (Daejung, 8587-4400), and then coverslipped with mounting medium (Sigma‒Aldrich, 03989).

To classify dendritic spines, the following established parameters and criteria were used. Dendritic spines were classified as mushroom, stubby, thin, or filopodium based on criteria from various studies^2–4^. Mushroom spines were identified by a head-to-neck diameter ratio (H/N) greater than 1.4 and a head diameter greater than 0.6 μm. Stubby spines were defined as those with no neck, a length-to-head ratio (L/H) less than 3, and a spine length less than 0.75 μm. After considering either mushroom or stubby spines, thin spines were defined as those with a spine length less than 3 μm, and filopodia were defined as those with a spine length greater than 3 μm.

Hippocampal slice preparation and electrophysiology

APP/PS1 mice and WT mice (13 months old) were anesthetized with isoflurane (5% isoflurane, 95% O_2_) and perfused with ice-cold sucrose artificial cerebrospinal fluid (aCSF) containing 195.5 mM sucrose, 2.5 mM KCl, 1 mM NaH2PO_4_, 32.5 mM NaHCO_3_, 11 mM glucose, 2 mM Na pyruvate, and 1 mM Na ascorbate (all chemicals from Sigma) bubbled with 95% O_2_/5% CO_2_ at a pH of 7.4. After perfusion, the brains were quickly removed from the skull, and sagittal hippocampal slices (400 μm thick) were cut on a vibratome (Leica). The slices were incubated at 35°C for 15 min in an incubation solution containing 119 mM NaCl, 2.5 mM KCl, 1 mM NaH2PO_4_, 26.2 mM NaHCO_3_, 11 mM glucose, 2 mM Na pyruvate, 1 mM Na ascorbate, 3 mM MgSO_4_, and 1.5 mM CaCl_2_. After incubation, the slices were transferred to aCSF solution at 23–24°C for 1 hour.

Field recordings were made with a concentric bipolar electrode positioned in the stratum radiatum of the CA1 region using an extracellular glass pipette (3–5 MΩ) filled with aCSF. Stimulation was delivered through a bipolar electrode (FHC, Bowdoin, ME, USA) placed in the SC-CA1 region. The SC circuit was visualized using differential interference contrast (DIC) microscopy at 4× magnification and identified by the ability to evoke short and constant latency fEPSPs at CA1 synapses by SC input stimulation. The test stimulation in all the fEPSP experiments was measured before the experiments (30–300 μA), and a test-pulse stimulation strength that evoked 50% of the maximum fEPSP was used. Baseline synaptic responses were recorded for 30 min, and then LTP was induced by theta burst stimulation (100 Hz, 10 trains, 40 ms duration, 200 ms intertrain interval). LTP was calculated by averaging the fEPSP amplitudes during the last 5 min of the recordings. Recordings were made every 10 s for 1 hour using an Axopatch 700A amplifier (Molecular Devices) digitized at 10 kHz and filtered at two kHz with Digidata 1440A and pClamp 10.0 software (Molecular Devices).

To measure mEPSCs, the electrode was filled with an internal solution containing 135 mM Cs methane sulfonate, 8 mM NaCl, 10 mM HEPES, 0.5 mM EGTA, 4 mM Mg-ATP, 0.3 mM Na-GTP, and 5 mM QX-315 Cl; pH 7.25 with CsOH, 285 mOsm). Miniature currents were recorded in the presence of tetrodotoxin (1 μM TTX, Tocris) to block sodium currents and propagate action potentials.

Stereotaxic FAM19A5 injection

For stereotaxic injection of WT and MT FAM19A5 (R58A, R59A), 3-month-old male and female mice were anesthetized with xylazine and ketamine. An injection cannula was stereotaxically inserted into the striatum (mediolateral, 2.0 mm from bregma; anteroposterior, 0.5 mm; dorsoventral, 3.5 mm) unilaterally (inserted into the right hemisphere). The infusion was performed at a rate of 0.2 μl/min, and 2 μl of FAM19A5 (diluted in PBS at a concentration of 5 μg/μl) or the same volume of PBS was injected into the mouse.

Animal behavioral tests

The male mice used in all the behavioral tests were 9–11 months old. All assays used littermates or age-matched animals. Behavioral tests were performed in a light- and noise-controlled behavioral room where the animals were allowed to adapt for one hour before each behavioral test. All the behavioral data were analyzed in a blinded manner.

Y-maze test

The Y-shaped maze consisted of three identical arms (40 cm in length, 15 cm in height) at a 120° angle from each other. Mice were allowed to freely explore the three arms for 8 min with a luminosity of 40 lux. The sequence and the total number of arms entered were measured. Spontaneous alteration (%) was calculated as follows: the number of triads containing entries into all three arms/maximum possible alternations (the total number of arms entered – 2) × 100. Mice that constituted fewer than 10 of the total arm entries were omitted from the analysis.

Morris water maze test

Mice were trained to find the hidden platform (9 cm in diameter) in a stainless-steel pool (90 cm in diameter, 50 cm in height) filled with water (22 ± 1°C) to a depth of 30 cm. The platform was 1 cm below the water level. In the acquisition phase, the mice were trained for six consecutive days with four trials/day. Each trial ended either when an animal climbed onto the platform or when a maximum of 60 s had elapsed. Next, the mice were allowed to remain on the platform for 10 s. If they failed to locate the platform within 60 s, they were guided to the platform and left there for 10 s, and the escape time was recorded as 60 s. On day 7, for the probe test, the platform was removed, and the mice were allowed to explore freely for 60 s. The spatial memory ability of the individual animals in the probe test was determined by the following parameters: latency to target, number of target crossings, distance to the target, percentage in the SW quadrant, and percentage in the target. The target and SW quadrants refer to the location and quadrant where the hidden platform was located, respectively. These parameters were analyzed by SMART video tracking software (Panlab). Mice displaying abnormal behaviors, such as freezing and floating, were excluded from the final data.

Passive avoidance test

The passive avoidance test, performed in plastic cages of identical size, assessed associative learning and memory by using the rodents’ innate preference for darkness. The inside of the darkened cage was designed to induce a foot shock; thus, when the mice entered the dark compartment from the light compartment, a foot shock (0.5 mA intensity) was applied for 3 s. To assess memory, we placed the mice in the light compartment 24 hours later and measured the time it took to enter the dark compartment. The maximal latency was 300 s. Animals that exhibited freezing behaviors (with an acquisition time ≥ 200 s) or abnormal exploration were excluded from the final analysis.

**Extended references**

1.	Kwak H, Cho EH, Cho EB, et al. Is FAM19A5 an adipokine? Peripheral FAM19A5 in wild-type, FAM19A5 knock-out, and LacZ knock-in mice 2020:2020.02.19.955351. doi:10.1101/2020.02.19.955351

2.	Tanaka T. Development of an efficient method for classifying large numbers of dendritic spines using confocal microscopy. *Juntendo Medical Journal.* 2021;67:329–32. doi:10.14789/jmj. JMJ21-R02

3.	Risher WC, Ustunkaya T, Alvarado JS, Eroglu C. Rapid Golgi Analysis Method for Efficient and Unbiased Classification of Dendritic Spines. *PLOS ONE.* 2014;9:e107591. doi:org/10.1371/journal.pone.0107591

4.	McGuier NS, Uys JD, Mulholland PJ. Chapter 9 - Neural Morphology and Addiction. In: Torregrossa M, editor. Neural Mechanisms of Addiction, Academic Press; 2019, p. 123–35. doi:10.1016/B978-0-12-812202-0.00009-9
